# Bone marrow B cell collapse promotes bone metastasis in breast cancer

**DOI:** 10.64898/2026.04.21.720007

**Authors:** Ana Teijeiro, Claudia A. Rivera, Motoyoshi Nagai, Adam X. Miranda, Eduard Ansaldo, Paula Juliana Perez-Chaparro, Bianca M. Nagata, Derron A. Alves, Niki M. Moutsopoulos, Yasmine Belkaid

## Abstract

Metastasis remains the primary cause of cancer-related deaths and is characterized by complex reprogramming of systemic processes. Emerging evidence indicates that extraosseous tumors can rewire bone marrow physiology and disrupt hematopoiesis, thereby compromising effective systemic immune responses. However, how tumor-induced immune alterations in bone marrow contribute to skeletal metastasis remains poorly defined. Here, using immunocompetent mouse models of mammary tumor bone metastasis, we show that mammary cancer cells precondition the bone marrow niche prior to metastatic colonization, driving early remodeling of the microenvironment and depleting bone marrow lymphoid populations. Specifically, cancer cells induce a dramatic B cell reduction, the most abundant lymphoid subset in bone marrow, resulting from dysregulated cell cycle gene expression in pre-B cells, along with impaired B-cell proliferation and differentiation. These findings are further validated in breast cancer bone metastasis patients, who exhibit significant bone marrow B-cell loss alongside disrupted molecular and developmental programs. A causal role for B cells in restraining skeletal metastasis is supported by the finding that experimental B-cell depletion significantly increases both incidence and severity of bone metastasis. Mechanistically, we find that B-cell loss is driven by systemic elevation of G-CSF. Accordingly, pharmacological neutralization of G-CSF significantly reduces both B-cell depletion and bone metastasis susceptibility. Collectively, our data reveal that breast cancer cells can distantly hijack B-cell developmental trajectories, promoting skeletal metastasis. This work identifies B cells and G-CSF as potential therapeutic targets in bone metastasis and highlights the importance of targeting early bone marrow immune dysregulation to prevent or limit skeletal metastasis.

**HIGHLIGHTS:**

- Mammary tumor cells reshape the bone marrow niche inducing B cell loss
- Bone marrow B cell development is impaired in mammary tumor metastasis
- Experimental depletion of B cells promotes bone metastasis
- G-CSF mediates B cell loss in mammary tumor metastasis

## INTRODUCTION

Metastasis remains the leading cause of cancer-related mortality and is a complex process involving the crosstalk of multiple organs that engage communication to maintain the body’s function.^1,2^

The establishment of metastasis depends not only on the intrinsic properties of cancer cells but also on the reprogramming of the immune, vascular, metabolic, microbiome, and neuroendocrine systems across the whole body.^3^ Therefore, metastasis needs to be understood and explored as a result of a systemic dysregulation rather than a localized process. More specifically, understanding the etiology of metastasis requires exploring how tumor cells hijack inter-organ communication networks.

The bone is a central regulator of fundamental physiological processes and it is also a common site of metastasis for many types of primary tumors, such as breast, prostate, lung, and kidney cancer.^4, 5^ Beyond its structural and endocrine functions, this organ serves as the primary reservoir of immune and progenitor cells, orchestrates hematopoiesis, and redistributes immune and progenitor cells in response to systemic signals.^6^ During metastasis, the bone becomes a critical target of systemic reprogramming. Increasing evidence revealed that extraosseous primary tumors can impose changes within the bone marrow prior to the occurrence of metastasis. For example, lung adenocarcinoma, melanoma, and breast cancers can disrupt normal bone function by promoting the proliferation, differentiation, and mobilization of bone marrow progenitor cells.^7–10^ These changes in the bone marrow immune dynamics create a systemic immune landscape that favors tumor growth and metastasis progression in distant organs. In the context of lung metastasis, mammary primary tumors skew hematopoiesis towards the myeloid lineage, leading to systemic expansion of granulocytes that in turn promote a pre-metastatic niche in the lungs.^9,11–13^ However, despite the growing recognition that altered hematopoiesis represents a point of vulnerability for the development of metastasis, the immune determinants that govern tumor colonization of the bone marrow remain poorly defined.

Breast cancer, the most common type of cancer worldwide, exhibits a particularly strong incidence of bone metastasis. Approximately 70% of patients with advanced breast cancer develop bone metastasis, making the bone the third most common metastatic location overall.^5^ Myeloid cells, osteolineage cells as well as T cell immunosuppression have been implicated in the establishment of pre-metastatic niches within the bone.^14–18^ However, despite its remarkable prevalence, the factors controlling breast cancer bone metastasis development remain poorly explored and understood. Unraveling how breast cancer progression perturbs the bone marrow immune microenvironment and how these alterations can support bone metastasis development is essential for developing strategies that target early components of disease progression.

In this study, we aimed to identify key immune factors that regulate bone marrow remodeling during mammary tumor metastasis. Our work uncovers bidirectional interactions between metastasis progression and bone marrow physiology and reveal a previously unappreciated role of B cells in the control of bone metastasis development.

## RESULTS

### Mammary tumor cells remodel the bone marrow lymphoid niche

To assess the impact of metastasis on the bone marrow immune microenvironment, we employed a well-established and relevant preclinical system associated with intracardiac injection (IC) of mammary tumor cells in immunocompetent mice **(Figure 1A)**. As previously described, this approach recapitulates the bone metastatic process, including cancer cell survival in the bloodstream, extravasation, and development of metastatic lesions in the bone marrow cavity.^19,20^ Of note, this administration route bypasses the initial steps of primary tumor growth, allowing for a focused investigation of the metastatic cascade. In the context of this work, we employed the E0771 cell line, derived from a C57BL/6 mouse spontaneous mammary gland tumor.^21,22^

**Figure 1.**
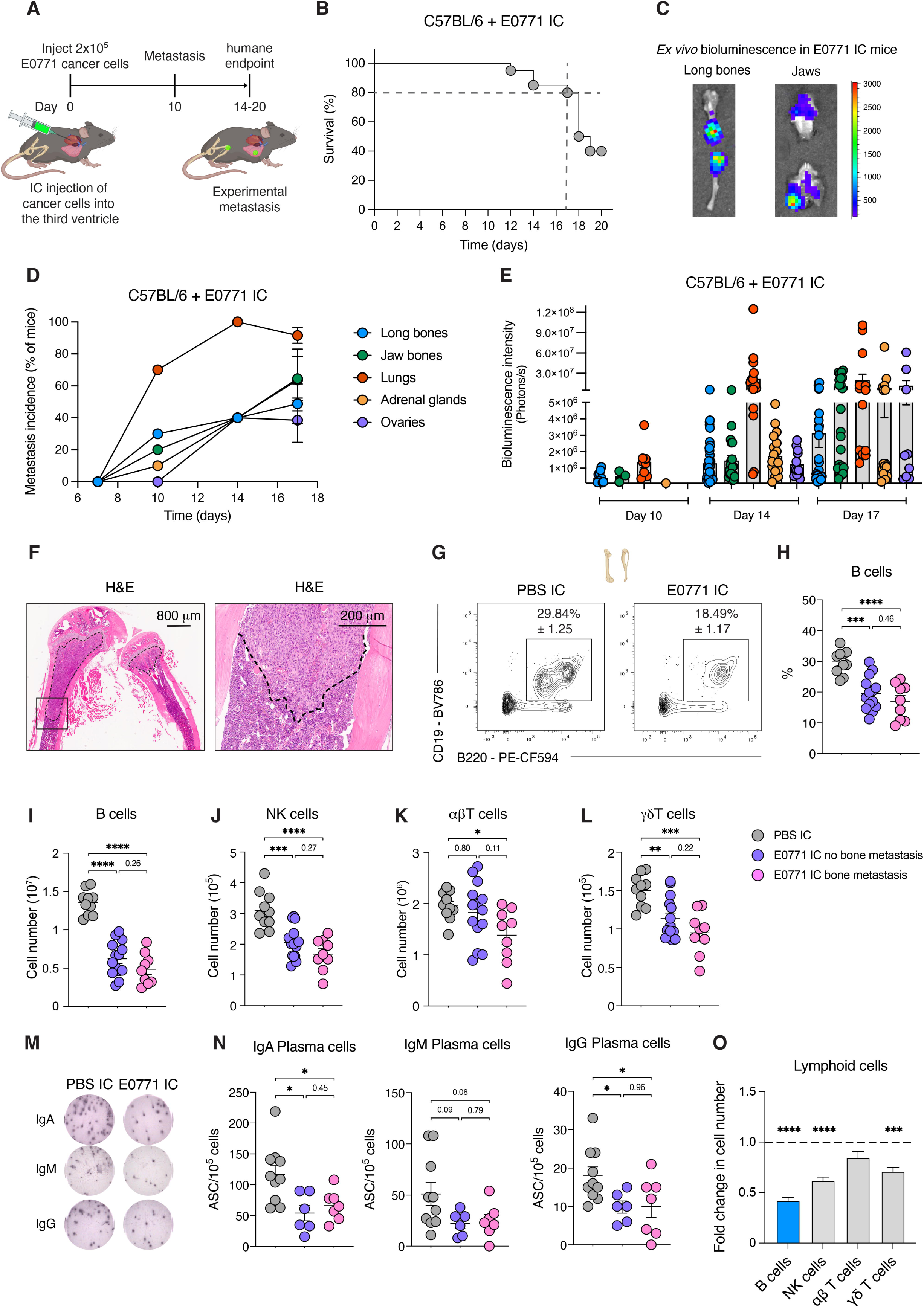
Mammary tumor cells remodel the bone marrow lymphoid niche. (A) Experimental layout. Intracardiac (IC) injection of E0771 mammary tumor cells is performed in C57BL/6 mice to induce experimental metastasis. (B) Survival curve of mice described in (A). (C) Representative images of *ex vivo* bioluminescence in long bones and jaws from mice described in (A). Color scale represents bioluminescence counts. (D, E) Metastasis incidence (D) and quantification of luciferase signal (E) in mice described in (A). Each dot represents an organ in (E). (F) Hematoxylin and eosin (H&E) staining of femur and tibia from E0771 IC mice showing experimental metastasis. Dashed line denotes marrow metastasis in a wedge-shaped pattern affecting the distal femur and proximal tibia and replacing bone marrow elements. (G) Representative contour plots of bone marrow CD45^+^ TCRβ^-^ CD138^-^ B220^+^ CD19^+^ B cells in C57BL/6 mice injected with PBS or E0771 cells IC at day 17. (H, I) Frequency (H) and absolute numbers (I) of bone marrow CD45^+^ TCRβ^-^ CD138^-^ B220^+^ CD19^+^ B cells in mice described in (G). (J-L) Absolute numbers of bone marrow NK cells (CD45^+^ CD90.2^+^ TCRβ^-^ NK1.1^+^) (J), αβT cells (CD45^+^ CD90.2^+^ TCRβ^+^ γδTCR^-^) (K), and γδT cells (CD45^+^ CD90.2^+^ γδTCR^+^ TCRβ^-^) (L) in mice described in (G). (M, N) Representative pictures of ELISPOT wells (M) and quantification (N) of the frequency of antibody secreting cells (ASCs) in bone marrow of mice described in (G). (O) Fold changes in absolute numbers of various lymphoid cells in bone marrow of E0771 IC mice with and without bone metastasis compared to PBS IC mice at day 17. Statistical analysis was performed using one-way ANOVA multiple-comparisons test (H-L and N) or unpaired two-tailed Student’s t test (O). Data are represented as means ± standard error of the mean (SEM). Each dot represents an organ in (E) and a mouse in (H-L and N). ∗p ≤ 0.05, ∗∗p ≤ 0.01, ∗∗∗p ≤ 0.001, ∗∗∗∗p ≤ 0.0001.

To precisely track neoplastic cell dissemination and disease progression via imaging techniques, E0771 cells were transduced to stably express GFP and luciferase using a lentivirus system **(Figure S1A).** IC injection of engineered E0771 cells into syngeneic C57BL/6 mice significantly impacted survival, with 60% lethality observed by day 20 **(Figure 1B)**. As such, we utilized day 17, a time point associated with 80% survival as the latest timepoint for future experiments. *Ex vivo* bioluminescence imaging revealed that mice developed progressive disease with disseminated cancer cells detected in various organs including bones, lungs, adrenal glands, and ovaries, as early as day 10 post-E0771 injection **(Figures 1C-1E and S1B)**. Of note, E0771 injected mice showed 40% and 60% incidence of metastasis in long bones and jaws, respectively **(Figure 1D)**. Histological analysis of long bones revealed that disseminated neoplastic cells filled the marrow cavity of the distal femur and proximal tibia, confirming bone metastasis **(Figure 1F)**. Microscopically, the wedge-shaped metastatic foci surrounded and efface bony trabeculae and replaced femoral and tibial bone marrow elements but generally do not infiltrate the growth plates of respective long bones **(Figure 1F)**. Thus, this approach represents a robust model system to explore the consequences of the potential interplay between bone marrow immune alterations and bone marrow metastasis.

We next performed a comprehensive profiling of the bone marrow immune landscape 17 days following injection with control (PBS IC mice) or E0771 cells (E0771 IC mice). While previous studies have examined how primary tumors reshape the myeloid lineage,^9–11^ the lymphoid compartment has received little attention. No significant differences were observed in the number of myeloid cells between E0771 injected and control mice **(Figure S1C)**. On the other hand, E0771 IC mice showed a profound and significant reduction in bone marrow lymphoid cells, including B cells, NK cells, αϕ3T cells, ψ8T cells, and plasma cells **(Figures 1G-1N)**. Of note, lymphoid cell loss was detected in bones of E0771 IC mice with or without bone metastasis. Thus, bone marrow immune alterations preceded the appearance of bone metastatic lesions. Among lymphoid cells, B cells, that represent the most abundant lymphoid cells within the bone marrow, were the most significantly impacted by cancer cells **(Figures 1O and S1D)**. To corroborate these findings, we profiled the bone marrow lymphoid landscape of another immunocompetent murine model of mammary tumor bone metastasis **(Figure S1E)**^23,24^. IC injection of 4T1.2 mammary tumor cells in syngeneic BALB/c mice also reduced the numbers of lymphoid cells, with a profound reduction in the B cell compartment **(Figures S1F-S1M)**. To determine if cancer cells may be able to impact the bone microenvironment prior to metastatic colonization, we analyzed mice bearing orthotopic (OT) mammary tumors (E0771 OT mice) **(Figure S1N)**. In line with previous studies^25^, E0771 cells failed to develop macrometastasis when orthotopically inoculated **(Figure S1O)**. Despite the absence of tumor cell dissemination, analysis of the bone marrow revealed marked alterations in the lymphoid landscape compared with tumor-free controls **(Figures S1P-S1U)**. Thus, mammary tumor cells can remotely influence the bone marrow prior to the presence of bone metastasis. Of particular interest was the marked reduction in B cells. This observation prompted us to investigate the mechanism underlying this collapse and the potential role of B cell depletion in tumor metastasis.

### Mammary tumor cells impair B cell development

Bone marrow is the primary site of lymphopoiesis. In particular, B cells arise from hematopoietic stem cells (HSCs) in the bone marrow and, once committed to the B cell lineage, progress through multiple developmental stages before migrating to secondary lymphoid organs, where they complete their maturation and enter circulation (**Figures 2A** and ^26^).

**Figure 2.**
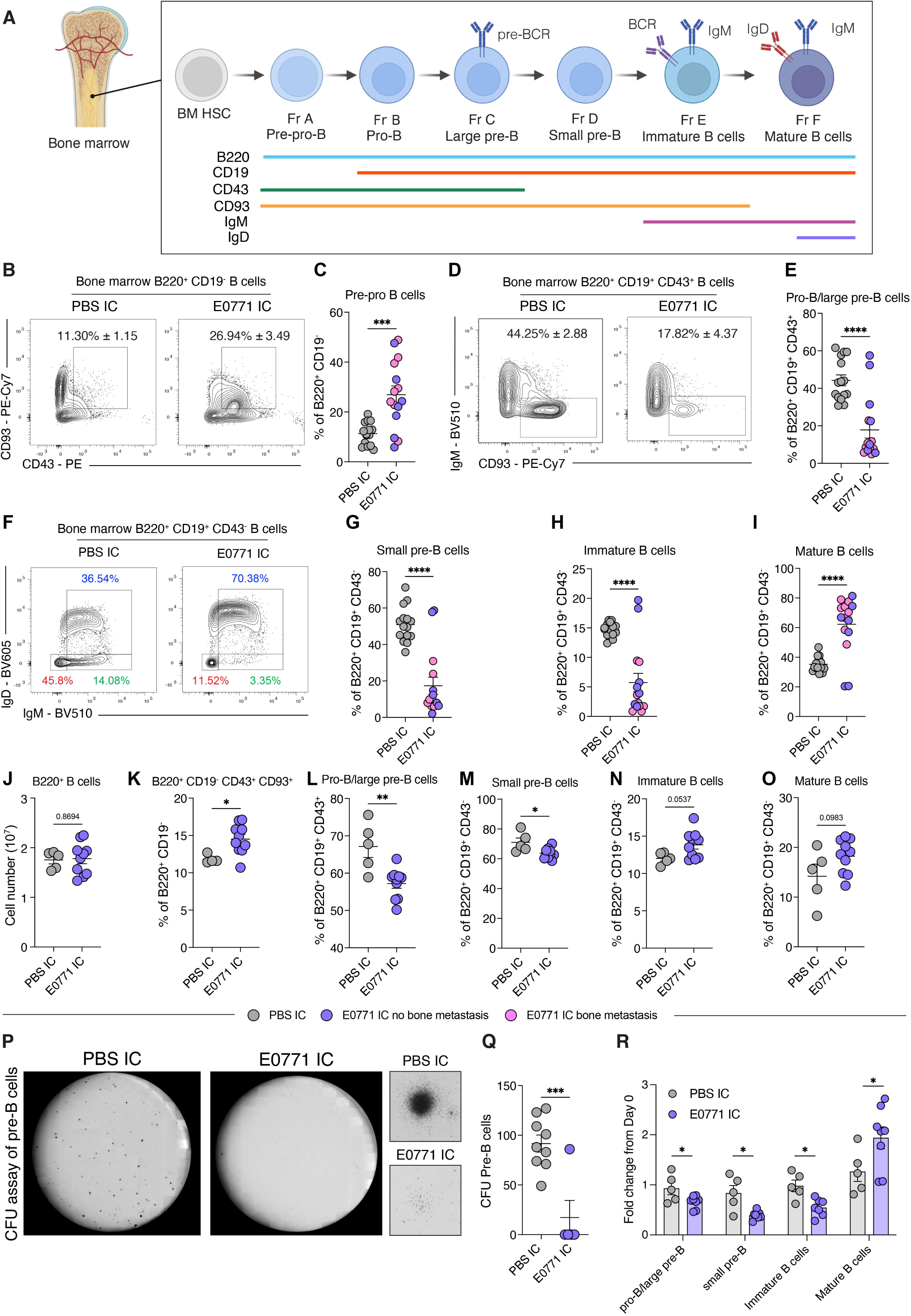
Mammary tumor cells impair B cell development. (A) Schematic representation of B cell lymphopoiesis and markers that define stages of B cell development. (B, C) Representative contour plots (B) and frequency (C) of bone marrow B220^+^ CD19^-^ CD43^+^ CD93^+^ cells in C57BL/6 mice with IC injection of PBS or E0771 cells at day 17. (D, E) Representative contour plots (D) and frequency (E) of bone marrow B220^+^ CD19^+^ CD43^+^ CD93^+^ IgM^-^ pro-B/large pre-B cells in mice described in (B). (F-I) Representative contour plots (F) and frequency of bone marrow B220^+^ CD19^+^ CD43^-^ IgM^-^IgD^-^ small pre-B cells (G), B220^+^ CD19^+^ CD43^-^ IgM^+^ IgD^-^ immature B cells (H), and B220^+^ CD19^+^ CD43^-^ IgM^+^ IgD^+^ mature B cells (I) in mice described in (B). (J-O) Absolute numbers of B220^+^ CD19^+^ B cells (J) and frequency of bone marrow B cell subsets (K-O) in C57BL/6 mice with IC injection of PBS or E0771 cells at day 7. (P, Q) Representative pictures (P) of colony formation unit (CFU) assay of pre-B cells and quantification of CFUs (Q) from bone marrow of mice described in (B). (R) Frequency of bone marrow B cell subsets from bone marrow of mice described in (B) cultured with IL-7 for 3 days, relative to day 0. Statistical analysis was performed using unpaired two-tailed Student’s t test (C, E, G-O, Q, R). Data are represented as means ± SEM. Each dot represents a mouse in (C, E, G-O, Q, R). ∗p ≤ 0.05, ∗∗p ≤ 0.01, ∗∗∗p ≤ 0.001, ∗∗∗∗p ≤ 0.0001.

We explored which B cell developmental stage was specifically impacted by tumor cells 17 days following IC injection of mammary tumor cells **(Figures 2A and S2A)**. We found that in addition of overall reduction, the relative representation of each developmental stage was significantly impacted compared to controls **(Figures 2B-2I and S2B-S2F)**. More specifically, mice harboring E0771 cells had significant increased proportions of B220^+^ CD19^-^ CD43^+^ CD93^+^ cells, that represent early B cell lineage-committed precursors **(Figures 2B and 2C)**. B220^+^ PDCA^+^ plasmacytoid dendritic cells (pDCs) were not increased **(Figures S2G and S2H)**, indicating that the decrease in B cells was not associated with a shift toward myeloid lineage differentiation. Among the B220^+^ CD19^+^ subset, the proportions of pro-B/large pre-B cells, small pre-B cells, and immature B cells were severely reduced **(Figures 2D-2H)**, while the proportions of mature B cells increased **(Figures 2F and 2I)**. Bone marrow B cells were also analyzed with the Hardy’s method ^27^, using BP-1/CD249 and HSA/CD24 to subset pro-B/large pre-B cells. This analysis revealed a reduction of Fractions B (pro-B cells) and C (large pre-B cells) in E0771 IC mice **(Figures S2I-S2L)**. Of note, the decrease in bone marrow B cells was not associated with significant changes in the absolute number of splenic B cells (B220^+^ CD19^+^) or splenic transitional B cells (B220^+^ CD19^+^ CD93^+^) **(Figures S2M-S2P)**. Collectively, these data support the idea that mammary tumor cells specifically impair B cell differentiation at the pro-B cell stage in the bone marrow.

To further dissect the sequence of events in the differentiation cascade impacted by mammary cancer cells, we next analyzed the bone marrow B cell landscape of E0771 injected at an early stage (day 7), a premetastatic time point at which no macrometastases were observed in any tissue (**Figure 1D**). Of note, changes in the proportions of B cell subsets were already detectable 7 days after IC injection of E0771 cells (**Figures 2J–2O**). Consistent with these observations, culture of bone marrow cells from E0771 IC mice in medium for pre-B cell colony-forming assay for one week showed a reduction in clonogenic B cell progenitors **(Figures 2P and 2Q)**. Further, *in vitro* differentiation of bone marrow B cells in IL-7 media showed that B cells from E0771 IC mice had reduced proportion of cells progressing to immature B cells **(Figure 2R)**. Thus, bone marrow B cell loss results from impaired B cell differentiation.

### Mammary tumor cells redefine the bone marrow B cell transcriptional landscape

To further characterize the cellular and transcriptional changes induced by mammary tumor cells in bone marrow B cells, we next performed single-cell RNA sequencing (scRNAseq) on sorted lymphoid cells from the bone marrow of PBS IC and E0771 IC mice **(Figure 3A)**. Clusters annotated as B cells were subset and classified by unsupervised hierarchical clustering and visualized with uniform manifold approximation and projection (UMAP). To characterize the B cell diversity, we used the ImmGen database and a reference-based annotation ^28,29^ **(Figures 3B, S3A, and S3B)**. In line with the flow cytometry analysis, E0771 IC mice showed significant decreased proportions of clusters identified as large pre-B cells (the transitional stage that proliferates significantly after expressing a functional heavy chain and pre-B cell receptor) **(Figures 3C and S3C)**.

**Figure 3.**
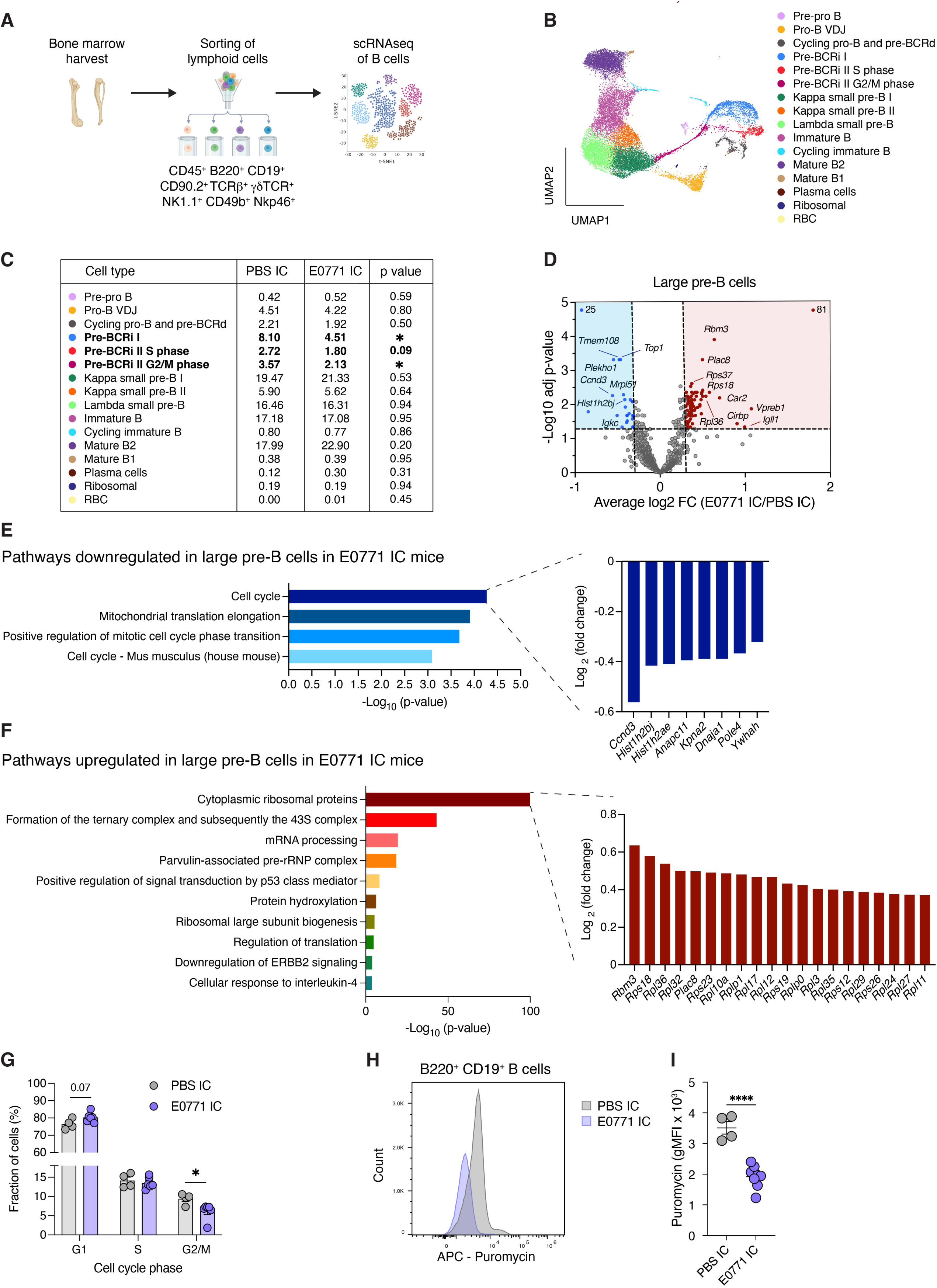
Mammary tumor cells redefine the bone marrow B cell transcriptional landscape. (A) Experimental design of single-cell RNA sequencing (scRNAseq). Lymphoid cells were sorted from bone marrow cells of C57BL/6 mice with IC injection of PBS or E0771 cells at day 17. Clusters identified as B cells were subset for analysis. (B) UMAP projection showing bone marrow B cell clusters analyzed by scRNAseq and annotated using previous reports.^29^ (C) Frequency of scRNAseq clusters of bone marrow B cells from mice described in (A). (D) Volcano plot displaying pseudobulk differentially expressed genes in large pre-B cells between PBS IC and E0771 IC mice. (E, F) Bar graph representing pathways downregulated (E) or upregulated (F) in large pre-B cells from E0771 IC mice relative to PBS IC mice. (G) Bar graph representing the percentage of cycling and non-cycling B cells from mice described in (A) analyzed by scRNAseq. (H, I) Representative histogram (H) and geometric mean fluorescence intensity (gMFI) (I) of puromycin incorporation in bone marrow B cells from mice described in (A). Statistical analysis was performed using unpaired two-tailed Student’s t test (C, G, I). Data are represented as means ± SEM. Each dot represents a cell in (B), a gene in (D), and a mouse in (G, I). ∗p ≤ 0.05, ∗∗∗∗p ≤ 0.0001.

To identify possible mechanisms responsible for the reduction of bone marrow B cells, we next characterized the transcriptional signature of B cells and large pre-B cells. Analysis of pseudobulk transcriptomes on biological replicates showed that IC injection of mammary tumor cells reprogrammed the transcriptomic landscape of B cells **(Figures 3D and S3D)**. Gene ontology pathway analysis of B cells revealed a downregulation of genes involved in cell cycle **(Figure S3E)**. Subsequent analysis of pre-B cells confirmed a downregulation of these pathways and was accompanied by a marked upregulation of ribosomal protein genes **(Figures 3E and 3F)**. These data proposed a shift toward a non-proliferative state in which cells reduce cell division while priming components of the protein synthesis machinery^30,31^. In line with these findings, analysis of the cell cycle status of B cell clusters revealed that IC injection of mammary tumor cells decreased the percentage of G2/M dividing bone marrow B cells **(Figure 3G)**. In agreement, B cells from E0771 IC mice showed decreased Ki67 expression **(Figures S3F and S3G)**. Further, despite the increase in genes encoding ribosomal proteins, B cells from mice injected with E0771 cells showed reduced puromycin incorporation **(Figures 3H and 3I)**, indicative of impaired translation. Taken together, these results show that the presence of mammary tumor cells impacts the transcriptional landscape of bone marrow B cells, impairing both cell proliferation and differentiation.

### Human breast cancer bone metastasis is associated with bone marrow B cell loss

To assess the clinical relevance of our findings, we took advantage of publicly available dataset and reanalyzed scRNAseq from bone marrow from healthy subjects and patients with pan-cancer bone metastasis, including breast cancer **(Figures 4A and 4B)**^32^. In line with our findings using experimental murine models, bone marrow samples from patients with breast cancer bone metastasis showed reduced B cell frequencies compared to healthy subjects **(Figure 4C)**. Furthermore, analysis of subset B cells revealed that the proportions of clusters of B cell progenitors also were reduced, while the clusters of mature B cells expanded, suggesting impaired differentiation **(Figures 4D and 4E)**. Next, we examined the pseudobulk transcriptomes of B cells. Importantly, gene ontology pathway analysis revealed that several genes involved in cell cycle processes were downregulated in B cells from breast cancer bone metastasis (**Figure 4F**). Additionally, analysis of the cell cycle stage showed that breast cancer bone metastasis significantly impaired B cell proliferation (**Figures 4G and 4H**). Taken together, these observations support the idea that B cell loss resulting from impaired proliferation and differentiation is also a characteristic of human bone metastasis and underscores the clinical relevance of our findings. The profound impact of mammary tumor cells on the B cell compartment in both experimental models and in the context of breast cancer patients prompted us to explore the possibility that this conserved response may contribute to the establishment of the bone premetastatic niche.

**Figure 4.**
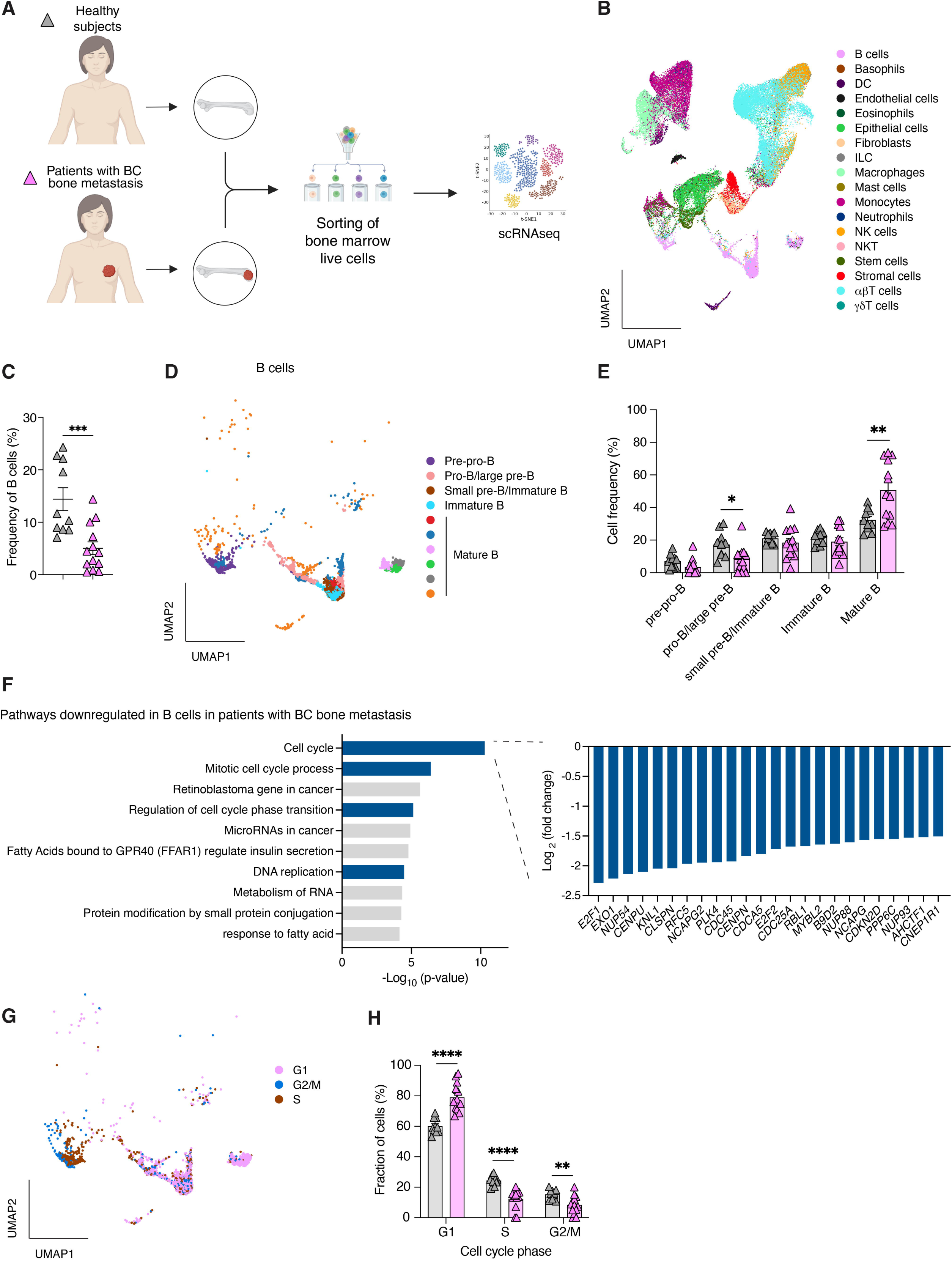
Human breast cancer bone metastasis is associated with bone marrow B cell loss. (A) Experimental design of scRNAseq as reported in ^32^. Live cells were sorted from bone marrow cells from healthy subjects or patients with breast cancer (BC) bone metastasis. (B)UMAP projection showing bone marrow cell clusters. (C) Frequency of bone marrow B cells from donors described in (A). (D) UMAP projection showing bone marrow B cell clusters analyzed by scRNAseq. (E) Frequency of scRNAseq clusters of bone marrow B cells from donors described in (A). (F) Bar graph representing pathways downregulated in B cells from donors described in (A). (G) UMAP projection representing cycling and non-cycling B cells from donors described in (A). (H) Bar graph representing the percentage of cycling and non-cycling B cells from donors described in (A) analyzed by scRNAseq. Statistical analysis was performed using unpaired two-tailed Student’s t test (C, E, H). Data are represented as means ± SEM. Each dot represents a cell in (B, D, G), and a subject in (C, E, H). ∗p ≤ 0.05, ∗∗p ≤ 0.01, ∗∗∗p ≤ 0.001, ∗∗∗∗p ≤ 0.0001.

### Experimental B cell loss promotes mammary tumor bone metastasis

Bone marrow B cell remodeling appears in early stages of the metastatic process, prior to the detection of macrometastasis in any organ **(Figures 2K-2O)**. This phenomenon supported the idea that loss of bone marrow B cells could create a vulnerable microenvironment prone to tumor colonization. Therefore, we next tested whether reduction of B cells could promote bone metastasis using genetically engineered mouse models of B cell depletion. We first used the μMT mouse model ^33^, which is congenitally deficient in B cells **(Figure 5A)**. Constitutive depletion of B cells in μMT mice significantly increased bone metastasis incidence as compared to wildtype mice **(Figure 5B)**. Moreover, bone metastatic burden was increased in long bones of μMT mice, as determined by higher total bioluminescence signal intensity **(Figures 5C and 5D)**. To corroborate these findings in a model allowing temporal depletion, we next used the *Cd19*-Cre-DTR inducible mouse model of B cell depletion^34^. Adult mice were treated with diphtheria toxin prior to the injection of E0771 cells (**Figure 5E**) to induce efficient depletion of B cells **(Figures S4A-S4G)**. Temporal depletion of B cells was associated with significant increase in bone metastasis incidence and severity compared to controls (**Figures 5F-5H**). Of note, the impact of constitutive or temporal B cell deficiency on metastasis was only observed in the bone marrow and not observed in lungs **(Figures S4H-S4K)**. Thus, experimental removal of B cells promotes specifically bone metastasis, supporting the idea that B cells directly or indirectly play a protective role.

**Figure 5.**
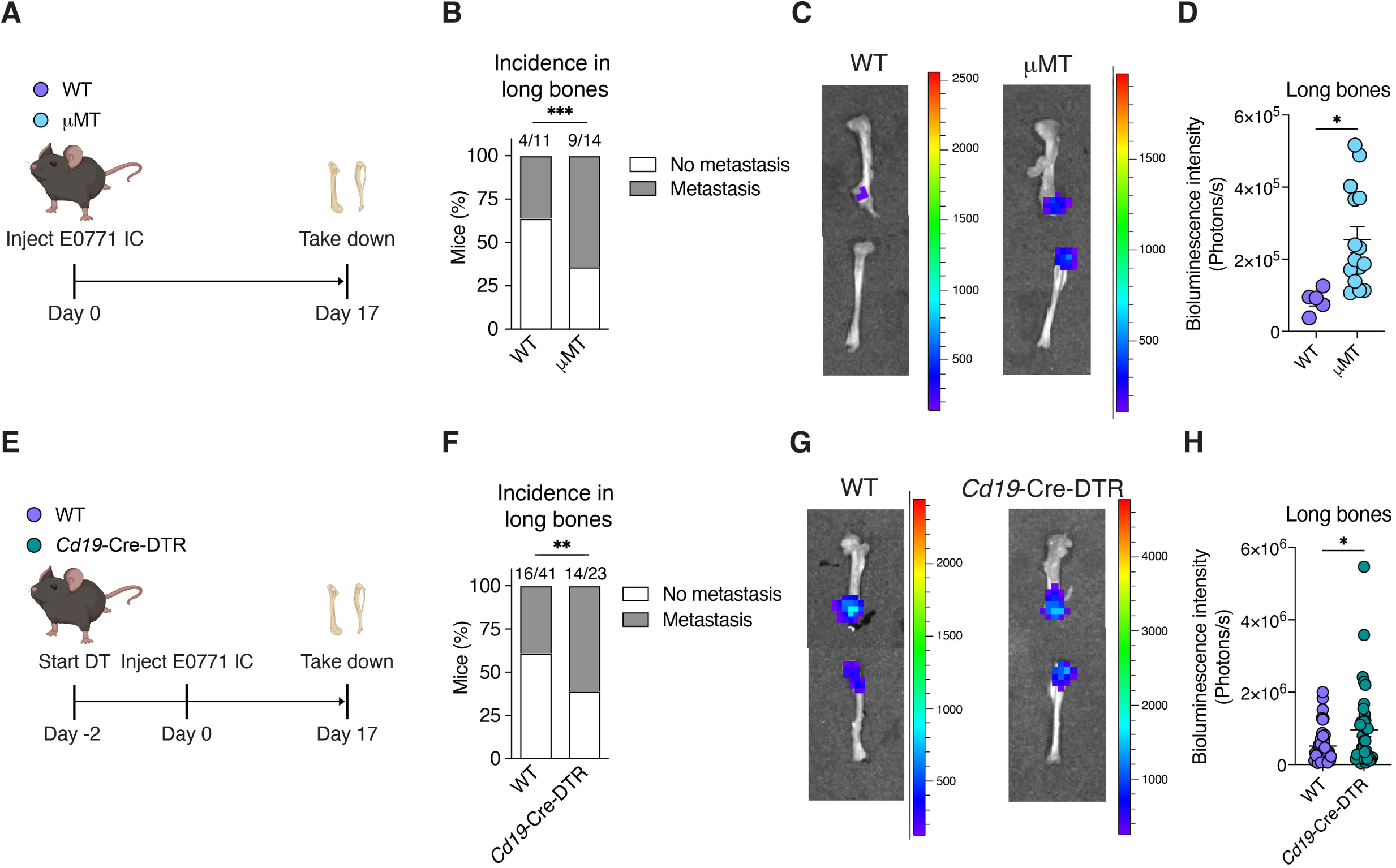
Experimental B cell loss promotes mammary tumor bone metastasis. (A) Experimental layout. Wildtype (WT) and μMT mice were injected IC with E0771 mammary tumor cells and analyzed 17 days after injection. (B) Bone metastasis incidence in mice described in (A). (C, D) Representative images (C) and quantification of luciferase signal (D) in femurs and tibias with mammary tumor metastasis in mice described in (A). Color scale represents bioluminescence counts. (E) Experimental layout. WT and *Cd19*-Cre-DTR mice were treated with diphtheria toxin (DT) and injected IC with E0771 mammary tumor cells. Mice were analyzed 17 days after IC injection. (F) Bone metastasis incidence in mice described in (E). (G, H) Representative images (G) and quantification of luciferase signal (H) in femurs and tibias with mammary tumor metastasis in mice described in (E). Color scale represents bioluminescence counts. Statistical analysis was performed using Fisher’s exact test (B, F) or unpaired two-tailed Student’s t test (D, H). Data are represented as means ± SEM. Each dot represents an organ in (D, H). ∗p ≤ 0.05, ∗∗p ≤ 0.01, ∗∗∗p ≤ 0.001.

### G-CSF is required for B cell loss and mammary tumor bone metastasis

Next, we investigated the potential mechanisms by which metastasis of mammary tumor cells could induce bone marrow B cell loss. Soluble factors, including cytokines, chemokines, growth factors, and metabolites can affect the bone marrow microenvironment and prepare the niche for tumor cell colonization.^35,36^ To explore a potential role for soluble factors, we measured the levels of cytokines and chemokines in the serum and bone marrow lysates of mice injected with E0771 cells. Differentially upregulated factors in E0771 IC mice in serum included granulocyte colony-stimulating factor (G-CSF), interleukin (IL)-6, IL-22, C-C chemokine ligand (CCL) 2/monocyte chemotactic protein 1, CCL7/monocyte chemotactic protein 3; while CXC motif chemokine ligand (CXCL) 5 and CXCL12 were downregulated **(Figure 6A)**. Changes in interferon gamma (IFN-γ),

**Figure 6.**
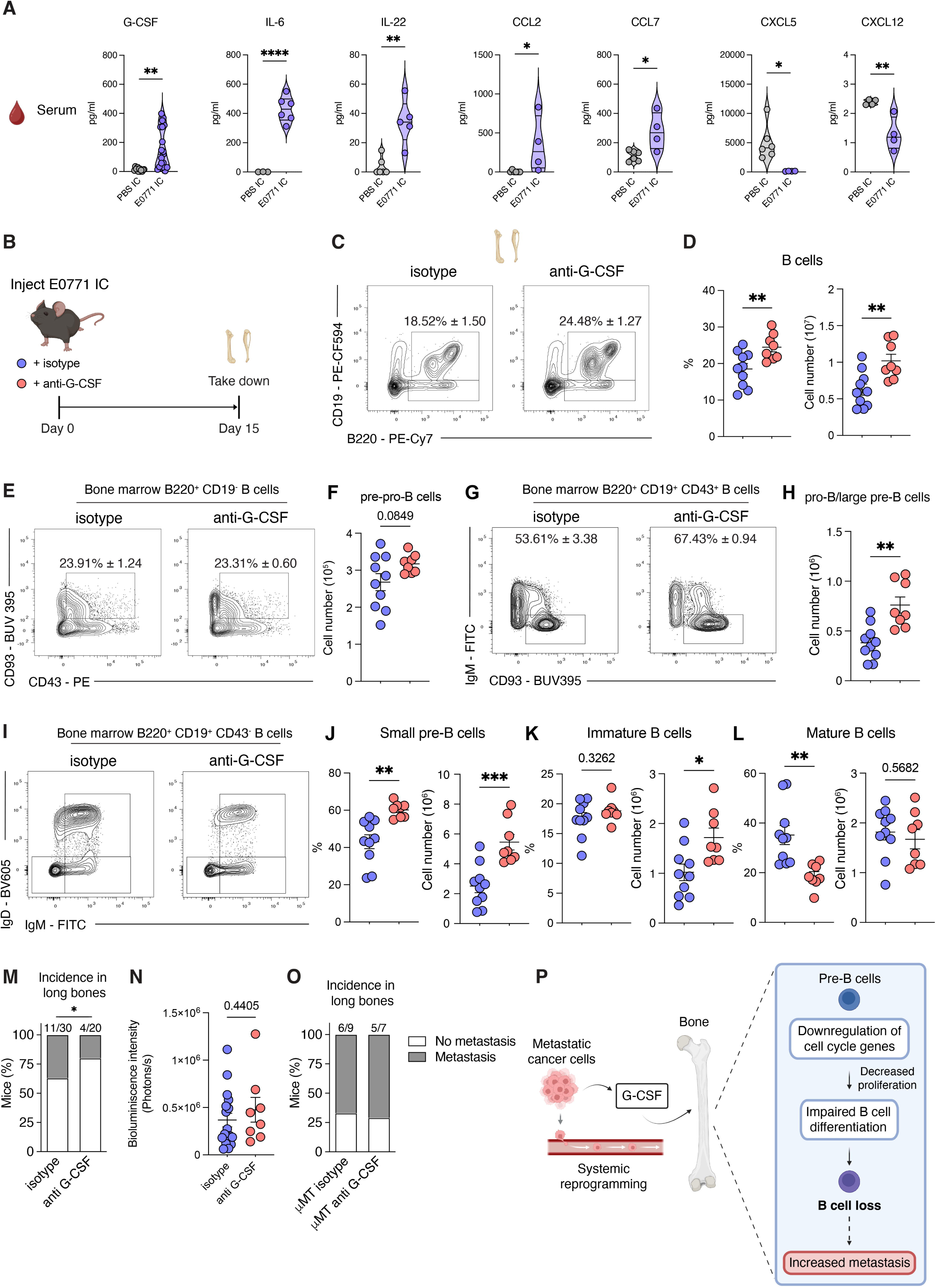
G-CSF is required for B cell loss and mammary tumor bone metastasis. (A) Cytokine and chemokine levels in serum of C57BL/6 mice with IC injection of PBS or E0771 cells at day 17. (B) Experimental layout. C57BL/6 mice were treated with G-CSF antibody or the isotype control and injected IC with E0771 mammary tumor cells. Mice were analyzed 15 days after IC injection. (C) Representative contour plots of bone marrow B220^+^ CD19^+^ B cells in mice described in (B). (D)Frequency (left) and absolute numbers (right) of bone marrow B220^+^ CD19^+^ B cells in mice described in (B). (E, F) Representative contour plots (E) and absolute numbers (F) of bone marrow B220^+^ CD19^-^CD43^+^ CD93^+^ cells in mice described in (B). (G, H) Representative contour plots (G) and absolute numbers (F) of bone marrow B220^+^ CD19^+^ CD43^+^ CD93^+^ IgM^-^ pro-B/large pre-B cells in mice described in (B). (I-L) Representative contour plots (I) and frequency of bone marrow B220^+^ CD19^+^ CD43^-^ IgM^-^IgD^-^ small pre-B cells (J), B220^+^ CD19^+^ CD43^-^ IgM^+^ IgD^-^ immature B cells (K), and B220^+^ CD19^+^ CD43^-^ IgM^+^ IgD^+^ mature B cells (L) in mice described in (B). (M, N) Bone metastasis incidence (M) and quantification of luciferase signal (N) in femurs and tibias with mammary tumor metastasis in mice described in (B). (O) Bone metastasis incidence in μMT mice treated with G-CSF antibody or the isotype control and injected IC with E0771 mammary tumor cells. Mice were analyzed 15 days after IC injection. (P) Schematic representation of the crosstalk between mammary tumor cells and the bone marrow to induce B cell loss and favor bone metastasis. Statistical analysis was performed using unpaired two-tailed Student’s t test (A, D, F, H, J-L, N) or Fisher’s exact test (M, O). Data are represented as means ± SEM. Each dot represents a mouse in (A, D, F, H, J-L) and an organ in (N). ∗p ≤ 0.05, ∗∗p ≤ 0.01, ∗∗∗p ≤ 0.001, ∗∗∗∗p ≤ 0.0001.

IL-17A, tumor necrosis factor alpha (TNF-α) or IL-10 were not detected (**Figure S5A**). On the other hand, the bone marrow niche only presented mild changes, with upregulation of CCL2, and CCL7 **(Figures S5B and S5C)**. Among the differentially detected factors, G-CSF was of specific interest, as G-CSF has been previously linked to suppression of B cell lymphopoiesis and reduction of pre-B cells^37,38^. Treatment of E0771 IC mice with G-CSF–neutralizing antibody significantly increased bone marrow B cell numbers (**Figures 6B–6D**). On the other hand, inhibition of IL-6, CCL2, CCL7 or CXCL12 signaling did not reverse B cell loss **(Figures S5D-S5H)**. G-CSF blockade also restored the proportions of bone marrow B cell subsets that were altered during mammary tumor metastasis (**Figures 6E–6L**). Notably, G-CSF neutralization was associated with an 17% reduction in bone metastasis incidence **(Figure 6M)**. Among mice that developed detectable lesions, photon flux values were similar between groups, suggesting that the G-CSF inhibition primarily limits metastatic establishment rather than metastatic growth **(Figure 6N).** G-CSF can directly promote neutrophil production^39^. However, at the low dose we employed, no differences in neutrophil numbers were observed (**Figures S5I and S5J**), supporting the idea that inhibition of G-CSF reduces bone metastasis by acting on the B cell compartment. To assess whether the impact of G-CSF on bone metastasis could be observed in mice devoid of B cells, μMT mice with IC injection of E0771 cells were treated with anti G-CSF. Notably, neutralization of G-CSF in mice lacking B cells had no impact on bone metastasis **(Figure 6O)**. Of particular interest, E0771 cells, similarly to other mammary tumor cells^40,41^, secrete G-CSF **(Figure S5K)**, supporting the idea that tumor cell-derived G-CSF could directly contribute to the B cell collapse. Taken together, these results support the idea that mammary tumor-derived G-CSF contributes at least in part, to bone marrow B cell loss, a phenomenon that in turn favors bone metastasis **(Figure 6P)**.

## DISCUSSION

Metastasis represents the most advanced and deadliest stage of cancer. Successful tumor cell colonization at distant sites requires a complex inter-organ communication and the remodeling of host physiology. How the bone marrow remodeling contributes to metastasis susceptibility remains poorly understood. Here, we showed that mammary tumor cells establish a crosstalk with the bone marrow at early stages of the metastatic disease, dramatically altering the bone marrow lymphoid microenvironment. Our work revealed that B cell loss occurs before macrometastasis is detected supporting the idea that this phenomenon contributes to the formation of the premetastatic niche in the bone and creates a permissive milieu for incoming tumor cells.

Our findings showed that mammary tumor metastasis causes a profound loss of bone marrow B cells. We showed that this effect was driven by impaired B cell differentiation due to altered expression of key cell cycle regulators and reduced proliferation of B cell progenitors. Notably, no differences were observed in the proportions of the pre-BCR-dependent proliferation stage, supporting the idea that the defect occurs after this checkpoint. B cell loss was restricted to the bone marrow and not observed within other organs such as spleen, suggesting that B cell loss was driven by bone marrow-specific regulatory mechanisms or systemic cues. Given that transitional and mature splenic B cells are long-lived cells,^42,43^ it is tempting to speculate that during mammary tumor metastasis, the spleen can sustain the B cell pool even without the influx of immature B cells.

Of particular importance, our data demonstrated that B cell loss is critical for metastasis permissiveness. Indeed, experimental depletion of B cells is sufficient to enhance bone metastasis susceptibility and severity. B cells are critical regulators of tissue physiology, contributing to immune functions through antibody production, antigen presentation, cytokine secretion, and interactions with immune and stromal cells.^44^ Whether the loss of B cells in the bone marrow creates a vulnerable niche through the alteration of unique regions of the bone marrow where B cells reside or via impairment of B cell functions remains an important and exciting area for future study. Depending on phenotype, spatial organization, and tumor type, B cells can exert opposing effects in tumorigenesis. While antigen-experienced B cells and plasma cells can promote anti-tumor immunity and correlate with better outcomes,^45–47^ IL-10/TGF-β-producing B regulatory cells (Bregs) can support tumor growth.^48^ In our model, IL-10 and TGF-β levels were unchanged, suggesting that Bregs may not drive bone metastasis. These context-dependent roles highlight the need to understand the role of B cells in metastasis to tailor targeted therapies that improve patient survival and quality of life. Our study broadens insights by showing that B cells are gatekeepers of mammary tumor bone metastasis.

Despite growing evidence implicating B cells in tumorigenesis, the tumor-derived and systemic cues governing their development and fitness remain unclear. Our results indicated that mammary tumor cells remotely alter bone marrow B cell differentiation and engrave unique molecular programs that support metastasis to the bone. Of note, we show that one of the factors contributing to B cell loss in mammary tumor bone metastasis is G-CSF. However, G-CSF neutralization did not completely abolish B cell loss, suggesting that additional pathways may contribute to the remodeling of the B cell compartment during tumorigenesis. G-CSF has pleiotropic functions, being essential for neutrophil production and hematopoietic stem cell mobilization into the bloodstream.^39^ Increasing evidence highlights the important role of G-CSF as a pro-metastatic factor in mammary tumor pulmonary metastasis through recruitment of Ly6C^+^Ly6G^+^ granulocytes and suppression of NK anti-tumor responses.^49–51^ Our work expands this knowledge by uncovering that G-CSF is critical in impairing B cell development and increasing bone metastasis susceptibility. In the bone marrow, G-CSF could act on numerous cell types.^52^ Previous studies suggest that G-CSF can impair B-lymphopoiesis in an extrinsic manner through its action on bone stromal cells.^53^ Whether G-CSF affects B cell development in an intrinsic or extrinsic manner in mammary tumor metastasis remains to be determined.

Of note, G-CSF treatment is used in cancer patients to improve chemotherapy-induced neutropenia and cell transplant ^51,52^. Since G-CSF administration also impairs B-lymphopoiesis in healthy donors,^54^ our work raises the important question of the role of G-CSF in human bone metastasis and suggests that, in patients with bone metastases, G-CSF exposure could paradoxically be associated with worse outcomes.

Bone metastasis is a common complication of cancer and is generally incurable. It causes considerable pain, bone fractures, and hypercalcemia. Unfortunately, therapeutic strategies to prevent or cure this disease are suboptimal. Bisphosphonates or receptor activator of NF-κB ligand (RANKL) antibodies are the standard of care for patients with bone metastases to reduce skeletal-related events.^5^ However, 30–50% of patients on these therapies still undergo disease progression.^55^ Our human data indicate that B cell loss also emerges in breast cancer bone metastasis, suggesting that B cells may also influence bone metastasis development in humans. Moving forward, it would be important to explore whether B cell-associated immunotherapies could be effective for patients with bone metastasis. Importantly, recent reports highlight that patients with bone metastasis have a poor response to immune checkpoint inhibitors (ICIs).^56,57^ This may be explained by the profound reduction of the lymphoid niche in the bone marrow, supporting the idea that these cells are critical players of the marrow physiology in metastasis. Further studies exploring how recovery of the homeostatic bone marrow lymphoid pool could boost the response to ICIs would be important to conduct.

Collectively, our work revealed B cells and G-CSF as key regulatory elements in the protection of the bone marrow during mammary tumor metastasis and further supports the notion of cancer as a systemic disease. Understanding how to precisely protect bone marrow homeostasis and more specifically the B cell compartment may open the door to novel therapies for patients with unmet needs.

## LIMITATIONS OF THE STUDY

The use of comprehensive mouse models that recapitulate the different metastatic stages is critical to understand the mechanisms underlying the disease. The E0771 IC is a relevant mouse model of mammary tumor bone metastasis that allows the use of most of the genetically engineered mice generated in C57BL/6, in contrast to the widely used 4T1.2 BALB/c mouse model. To further validate our findings, the development of a spontaneous bone metastasis model in immunocompetent mice that more closely recapitulates the human disease will be necessary in the future. Furthermore, while we have identified B cells as critical mediators of bone metastasis development, we cannot dissect the specific role of bone marrow B cells and systemic B cells, since our models deplete B cells systemically. The generation of molecular markers that define tissue-specific B cells and genetic tools that allow specific targeting of B cell subsets will be critical for a comprehensive understanding of the role of B cells in bone metastasis. Finally, this study focused on the upstream signals that regulate B cell loss in mammary tumor metastasis. However, the downstream mechanisms by which B cells protect from bone metastases require further investigation.

## RESOURCE AVAILAVILITY

### Lead contact

Further information and requests for resources and reagents should be directed to and will be fulfilled by the lead contact, Yasmine Belkaid and Ana Teijeiro.

### Materials availability

Reagents generated in this study are available from the lead contact with a completed materials transfer agreement.

### Data and code availability

scRNA-seq data have been deposited into the Gene Expression Omnibis (GEO) data repository with the accession number GSE298523. All data are available in the main text or the supplementary materials.

## ACKNOWLEDGEMENTS

This research was supported by the Intramural Research Program of the National Institutes of Health (NIH). The contributions of the NIH authors are considered Works of the United States Government. The findings and conclusions presented in this paper are those of the authors and do not necessarily reflect the views of the NIH or the U.S. Department of Health and Human Services. This work was supported by the NIAID Division of Intramural Research. A.T., M. N., C.A.R., E.A., are supported by the Division of Intramural Research of NIAID (NIAID; 1ZIA-AI001115 and 1ZIA-AI001132); A.T. is an EMBO Postdoctoral Fellow (ALTF 1014-2021); C.A.R. is the Lorraine W. Egan Fellow of the Damon Runyon Cancer Research Foundation (grant DRG-2496-23); M.N. is a JSPS Research Fellow; Y.B. is supported by Institut Pasteur. We thank E. Bewaji, K. Webber, and NIAID animal facilities for technical support; Histoserv, Inc technical staff for preparing mouse samples for histopathological analysis; M. Smelkinson for assisting with imaging assays; P. Schwartzberg, E. Segrist, and N. Bouladoux for critical reading of the manuscript; and all the members of the Belkaid and Moutsopoulos Laboratories for helpful discussions and for providing constructive feedback on this project. Schematic figures were created with BioRender.

## AUTHOR CONTRIBUTIONS

A.T. and Y.B. conceived, designed the study and experiments, and wrote the manuscript. A.T. performed the experiments, analyzed the data, and created the figures. C.A.R., M.N., and P.J.P.C participated in performing experiments. A.X.M. and E.A. assisted with scRNA-seq analysis. B.M.N. assisted with the preparation of mouse samples for histopathological analysis. D.A.A. performed histopathology of mouse tissues. N.M.M. provided intellectual expertise and shared key methodologies. All the authors reviewed and contributed to editing the manuscript before submission.

## METHODS

### Cell lines

E0771 (CRL-3461) and 4T1.2 (CRL-3406) mouse mammary tumor cells were obtained from ATCC. E0771 cells were cultured in DMEM (Corning, 10-017-CV) supplemented with 20 mM HEPES (Corning, 25-060-Cl), 10% fetal bovine serum (FBS), 100 unit/ml penicillin and 0.1 mg/ml streptomycin (1 × P/S). 4T1.2 cells were cultured in alpha-MEM (Gibco, 12561-056) supplemented with 10% FBS and 1 × P/S. E0771 cells were engineered to stably express the CMV-Luc-GFP-puro luciferase and fluorescence dual reporter (LipExoGen Biotech, LFV-0001-1S) according to the manufacturer’s instructions. Cells expressing the dual reporter were selected with puromycin and fluorescence-activated cell sorting (FACS) of GFP positive cells.

### Mice

Specific pathogen-free (SPF) C57BL/6 CD45.1.2 (C57BL/6J x B6.SJL-CD45a(Ly5a)/Nai F1 (Taconic line 8422), μMT mice (*Ighm^tm1Cgn^*, Taconic line 49),^33^ and CCR2 knockout mice (*Ccr2^tm1Ifc^*, Taconic line 8456)^58^ were obtained from the NIAID Taconic Exchange Program. *Cd19*-Cre mice (*Cd19^tm1(cre)Cgn^*)^59^ and ROSA26iDTR mice (*Gt(ROSA)26Sor^tm1(HBEGF)Awai^*)^60^ were obtained from the Jackson Laboratory and crossed to obtain *Cd19*-Cre-DTR mice. All mice were bred and maintained at an American Association for the Accreditation of Laboratory Animal Care (AAALAC)–accredited animal facility at NIAID and housed in accordance with procedures outlined in the *Guide for the Care and Use of Laboratory Animals*. All experiments were performed at NIAID under an animal study proposal (LHIM2E and LHIM3E) approved by the NIAID Animal Care and Use Committee. Only female mice were used for this study. Mice were fed a chow diet (Lab Diet Advanced Protocol, 5V75). Food and water were provided *ad libitum*. Age-matched 8-12 weeks of age mice were used in each experiment. Littermate controls were used for experiments involving mice bred and maintained at NIAID. Mice were randomly assigned to either experimental or control groups. Mouse samples were harvested from 7am to 12pm.

### Mouse *in vivo* treatments

For inducible B cell depletion, diphtheria toxin (DT) (List labs, #150) was dissolved in PBS and 10 ng/g (body weight) was intraperitoneally injected. Mice were treated 2 consecutive days before injection of tumor cells and then 3 times per week until the completion of the study. For G-CSF blocking, mice were intraperitoneally injected with 25 μg of Rat IgG2a isotype control or anti-mouse G-CSF (Invitrogen, 16-7353-85) every other day. For IL-6 neutralization, mice were intraperitoneally injected with 200 μg of Rat IgG1 isotype control or anti-mouse IL-6 (BioXcell, BE0046) every other day. For CXCR4 inhibition, PBS vehicle or 10 mg/kg AMD3100 (Abcam, AB120718) were injected daily. For CCL7/MCP-3 neutralization, mice were injected with 20 μg of mouse CCL7/MCP-3 antibody (R&D systems, AF-456-NA) or control IgG antibody every 3 days.

### Induction of experimental mammary tumor metastasis

Animals were anesthetized with 2% isoflurane and 2 x 10^5^ E0771 cells or 1 x 10^5^ 4T1.2 cells resuspended in 100 μL of PBS were injected into the left cardiac ventricle using a 31 G syringe needle (BD, 328438). Mice were weighed and monitored daily for respiratory distress, decreased activity, ataxic gait, or signs of paralysis for up to 20 days after injection and euthanized at the indicated time points or when they reached humane endpoints. Mice with visible extrapleural intrathoracic tumors were excluded from the analysis. Metastasis incidence was determined by *ex vivo* bioluminescence imaging, and subsequent histopathological analysis by H&E or flow cytometry analysis using markers expressed by mammary tumor cells. Bone marrow cells from bones processed for flow cytometry analysis were cultured in media to grow mammary tumor cells for 1 week to confirm or discard the presence of neoplastic cells.

### Induction of orthotopic mammary tumors

2 x 10^5^ E0771 cells resuspended in 100 μL of PBS were injected into the 4^th^ mammary fat pad using a 31 G syringe needle (BD, 328438) to generate orthotopic mammary tumors. Mice were weighed daily and tumor volume was measured every two days with a digital caliper. Tumor size was calculated by measuring the length and width of the tumor and using the formula: (width)^2^(length)/2.

### Ex vivo bioluminescence imaging

Mice were retro-orbital injected with 150 mg/kg of D-luciferin 5 min before being euthanized by cervical dislocation. Tissues were quickly removed and washed with RPMI media. Bioluminescence images of the organs were acquired with IVIS BLI system (PerkinElmer, Caliper Life Sciences) with the following conditions: open emission filter, exposure time = 1 min, binning = large: 8, field of view = 12.5 ×12.5 cm, and f/stop = 1. Images were analyzed with Aura software. After imaging, samples were freshly processed for flow-cytometry analysis, processed in formalin for paraffin embedding, or snap frozen and kept at –80 °C until further analyses.

### Tissue processing for flow cytometry

To isolate cells from the bone marrow, femurs and tibias from one hind limb were flushed with RPMI media, filtered using a 70-μm nylon cell strainer, and centrifuged at 500g for 5 min at 4 °C. To isolate cells from the spleen, spleens were minced, dissociated and filtered through a 70-μm nylon cell strainer and centrifuged at 500g for 10 min at 4 °C. After centrifugation, supernatant was discarded and pellet was resuspended in PBS buffer for flow cytometry analysis.

### Flow cytometry analysis

Single cell suspensions were incubated with LIVE/DEAD Fixable Blue Dead Cell Stain Kit (Invitrogen Life Technologies) for 30 min at 4°C to discriminate live/dead cells. After washing the cells with PBS, cells were stained with fluorochrome-conjugated antibodies against surface markers in PBS for 30 min at 4 °C. After surface staining, cells were fixed and permeabilized using the Foxp3/Transcription Factor Staining Buffer Set (ThermoFisher Scientific) for 45 min at 4°C and stained with fluorophore-conjugated intracellular antibodies for at least 1 hour at 4°C. Stainings were performed in the presence of purified anti-mouse CD16/32 (Fc block). After washing, cells were resuspended in FACS buffer and analysed using LSR Fortessa (BD BioSciences). Flow cytometric data was analyzed using FlowJo software (Tree Star).

### Tissue processing for H&E

The Infectious Disease Pathogenesis Section at the National Institutes of Health or a commercial service (Histoserv, Inc., Germantown, MD) was utilized for tissue processing. Femurs and tibias were harvested, fixed in 10% neutral buffered formalin, decalcified, and embedded in paraffin according to the standardized protocols at the respective facilities. Select non-bone tissues (e.g., lung, heart, and mammary adipose tissue) were also routinely collected and processed. Samples were sectioned at 3-5 µm and stained with hematoxylin-eosin (H&E) for histopathological assessment. Samples were evaluated by a board-certified veterinary pathologist in a blinded manner. Sections were examined under light microscopy using an Olympus BX51 microscope and photographs were taken using an Olympus DP28 camera. Histopathological assessment of experimental metastasis in long bones was based on routine H&E staining.

### ELISPOTS

For the ELISPOT assay, Elispot 96-well plates with hydrophobic PVDF membrane (Millipore, MSIPN4W50) were coated with 5 μg/ml goat anti-mouse immunoglobulins (Southernbiotech: IgM human ads-UNLB, 1020-01; IgG human ads-UNLB, 1030-01; IgA human ads-UNLB, 1040-01) overnight at 4 °C in a humidified chamber. Wells were washed 4 times with PBS and after blocking with 0.1% BSA in PBS, 1.5 x 10^5^ bone marrow cells resuspended in RPMI media with 10% FBS were incubated for 3 hours in a humidified incubator at 37 °C and 5% CO_2_. After the cells were washed 4 times with PBS, alkaline phosphatase-conjugated goat anti-mouse antibodies (Southernbiotech: IgM-AP, 1020-04; IgG-AP, 1030-04; IgA-AP 1040-04) in 0.1% BSA in PBS were incubated overnight at 4 °C in a humidified chamber. After a wash, wells were incubated with alkaline phosphatase substrate BCIP/NBT (Sigmafast, B5655) to visualize antibody secreting cells (ASCs) marked with the bound antibodies. Reaction was stopped with tap water and ASCs were counted using an ImmunoSpot analyser (CTL) with CTL Switchboard 2.6.1.

### *In vitro* pre-B cell colony-forming unit (CFU) assay

Femurs and tibias from one hind limb were flushed with PBS with 2% FBS, filtered using a 70-μm nylon cell strainer, and centrifuged at 500g for 5 min at 4 °C. After centrifugation at 500g for 10 min at 4 °C, supernatant was discarded and pellet was resuspended in 1 ml of ACK lysing buffer (Quality Biological, 118-156-721) for 5 min at room temperature (RT) to lyse red blood cells. The reaction was stopped by adding 9 ml of PBS. After centrifugation at 500g for 10 min at 4 °C, cells were resuspended in PBS with 2% FBS. CFU assays were performed with methylcellulose media for pre-B lymphoid progenitor cells (STEMCELL Technologies, MethoCult^TM^ M3630 media) according to the manufacturer’s instructions. Briefly, 2.5 x 10^5^ cells were diluted in 2.5 ml of methocult media and dispensed as duplicates (1ml per well with 10^5^ cells) in 6-well cell culture plates (Cellstar, 657 160). Plates were incubated in a humidified incubator at 37 °C and 5% CO_2_ for 7 days. Pre-B cell colonies per well were evaluated and counted from images taken with an inverted microscope.

### *In vitro* culture of B cells

Bone marrow was isolated from femurs and tibias and depleted from erythrocytes using ACK lysing buffer as described above. 7.5 x 10^5^ cells/ml were cultured in IMDM media containing Glutamax (Gibco, 31980030) supplemented with 10% FBS, 1 × P/S and 5μM 2-Mercaptoethanol, and 10 ng/ml of recombinant mouse IL-7 (Invitrogen, RP-8664) in a humidified incubator at 37 °C and 5% CO_2_.^61^ Cells were harvested 3 days later for phenotypic characterization of B cells by flow cytometry analysis. Proportions of different B cell subsets were normalized to the values from day 0.

### Single-cell RNA sequencing analysis of mouse bone marrow cells

1.5x10^4^ bone marrow lymphoid cells (CD45^+^ B220^+^ CD19^+^ CD90.2^+^ TCRβ^+^ γδTCR^+^ NK1.1^+^ CD49b^+^ Nkp46^+^) from 5 PBS IC mice and 7 E0771 IC mice (5 mice with bone metastasis and 2 mice without bone metastasis) were sorted from samples labeled with TotalSeqC hashtags antibodies (Biolegend) using a Sony MA900 sorter. Sorted cells were pooled during sorting, and loaded on a Chromium Single Cell Controller (10X Genomics) to encapsulate the cells into droplets. The gene expression (GEX) and hashtag oligonucleotide (HTO) sequencing libraries were prepared using Chromium Single Cell 5’ v2 reagent kits following the manufacturer’s instructions and sequenced on an Illumina Nextseq2000 platform (NextSeq 1000/2000 P2 reagents). The initial processing of the expression data involved generating fastq files and count matrices using Cell Ranger v7.1.0^62^ (10X Genomics, Pleasanton, CA) that was run with the “Include introns=True” option and the mm10-2020-A transcriptome reference. Processing of the expression data was performed with the ‘cellranger count’ and ‘cellranger aggr’ functions.

After processing and aggregation of the lymphoid data, the estimated number of cells and mean reads per cell were as follows: 45127 estimated number of cells, 24305 mean reads per cell (from 3 GEX libraries), 535 Median hashtag UMIs per Cell (across 3 HTO libraries). The overall sequencing quality was high in the 3 mRNA libraries: >93.3% of bases in the RNA reads had Q30 or above. Furthermore, across the 3 GEX libraries the median gene count per cell range was 1508-1735, the sequencing saturation ranged between 68-72%, and 73-78% of the reads mapped confidently to the transcriptome.

The downstream analysis of the lymphoid data was performed in 4.4.2, the Cell Ranger aggregation output (filtered_feature_bc_matrix directory) was loaded into Seurat version 5.1.0^63^ using the CreateSeuratObject function with min.cells=3. For filtering out low-quality cells, we established thresholds for gene and UMI counts (nCount_RNA > 1000, nFeature_RNA > 300), and a maximum threshold for percentage mitochondrial content (percent.mt < 6) based on visual inspection of the corresponding metric distributions across all cells. Log-normalization (LogNormalize) of the RNA count data and centered log-ratio transformation (CLR) of the HTO count data were carried out using the NormalizeData function. For HTO demultiplexing cells into the individual mice and removing any doublets, HTODemux was run with a positive quantile = 0.99 and only cells identified as Singlets were kept. The top 2000 variable features were determined with the VariableFeatures function, expression data was scaled with ScaleData, and dimensionality reduction was performed with the RunPCA function on the variable features, followed by determining the dimensionality of the dataset on an elbow plot (ElbowPlot function), which was determined to be 50. Subsequently, FindNeighbors, FindClusters, and RunUMAP were run on 50 dimensions and a range of resolutions to determine the optimal resolution that captures biological diversity in the dataset, which was determined to be 0.4, resulting in 19 clusters. The clusters identified by the FindClusters function of Seurat (default Louvain clustering setting and 0.4 resolution) were annotated using cell type specific markers, cluster specific differentially expressed genes, and with the aid of SingleR,^64^ using the ImmGen database reference.^28^ One sample (‘B6-PBS-1’) was omitted from further analysis because it was identified as an outlier.

For analysis of B cells and B cell precursors, clusters annotated as B cells or B cell precursors were subset and FindVariableFeatures (nFeatures=2000) was run, data was scaled with ScaleData, RunPCA was run (npcs=100) and the top 28 dimensions were used to run FindNeighbors, FindClusters (with a range of resolutions and a final chosen resolution of 0.43), and RunUMAP. The resulting clusters were annotated using cell type specific markers, cluster specific differentially expressed genes, a reference-based annotation,^29^ and with the aid of SingleR,^64^ using the ImmGen database reference.^28^ pDCs were excluded from this analysis.

For pseudobulk analysis, the raw counts were aggregated across cells for each biological replicate on the indicated clusters using AggregateExpression using Seurat 5.1.0^63^ in R 4.4.2. Genes with a very low level of expression across biological replicates, less than 10 total counts across all samples, were filtered from the dataset. Differential gene expression analysis was performed using DESeq2 v1.44.0^65^ on the aggregated raw counts, using default parameters, and not including a log2 fold change shrinkage step. The normalized counts were extracted from the DESeq2 object and used for visualization along be corresponding adjusted p-values and log2 fold changes. A log2FC threshold of 0.3 and an adjusted p value threshold of 0.05 were used. Pathway enrichment analysis of upregulated or downregulated genes was performed using Metascape.^66^

For the analysis of the cell cycle phase, cells were scored for expression of cell cycle genes using the CellCycleScoring function of Seurat 5.1.0.^63^ The cell cycle markers used by Seurat are defined by the Kowalczyk et al 2015 study.^67^ This function classifies cells into S, G1, and G2/M cell cycle phases by the expression of those cell cycle markers.

### Puromycin incorporation

Bone marrow cells were isolated as described before. 2 x 10^6^ cells in RPMI media were incubated with 10 μg/ml of puromycin (RRID: AB_2827926) for 45 min in a humidified incubator at 37 °C and 5% CO_2_. After puromycin treatment, cells were washed in cold PBS and cell suspensions were incubated with fluorochrome-conjugated antibodies as described above and analyzed by flow cytometry.

### Measurement of cytokines and chemokines

Blood was collected from the heart and transferred to a serum tube with clot activators. Serum was collected after 4,000g, 20-min centrifugation at 4 °C, and kept at −80 °C until further analyses. Frozen femurs and tibias were lysed using RIPA lysis buffer (Cell Signaling, 9806) containing 2mM EDTA (Corning, 46-034-Cl), 0.5 mM PMSF and a protease/phosphatase inhibitor cocktail (Cell signaling, 5872). Cytokines and chemokines were quantified using the ProcartaPlex™ Mouse Cytokine & Chemokine Panel 1A, 36plex (Invitrogen, EPX360-26092-901), following manufacturer’s instructions. Differentially expressed cytokines and chemokines were confirmed by ELISA for the protein of interest. G-CSF levels were measured with Mouse G-CSF (CSF3) ELISA Kit (Thermofisher, EMCSF3). IL-6 levels were measured with Mouse IL-6 Quantikine ELISA kit (R&D Systems, M6000B). CXCL12 levels were measured with the Mouse CXCL12/SDF-1 alpha Quantikine ELISA Kit (R&D Systems, MCX120). CCL7/MCP-3 levels were measured with the Mouse MCP-3 Instant ELISA™ Kit (Invitrogen, BMS6006INST).

### Single-cell RNAseq analysis of human bone marrow samples

Publicly available single-cell RNA-seq data were obtained from Lui et al., 2025^32^. The dataset is available at the NCBI GEO database under accession number GSE266330. Data was accessed from the linked Zenodo archive (https://zenodo.org/records/16937964) on December 8, 2025. Breast cancer bone metastasis and control samples were selected for analysis from the “47.integrated.rds” file. Cell type prediction of the selected scRNA samples was carried out using the ImmGen database (Version: 2024-02-26) from the celldex package^64^ (Version: 1.16.0) and the SingleR package^64^ (Version: 2.8.0). To perform this cell type prediction, the human gene IDs of the single cell samples needed to be converted to their mouse ortholog counterparts using the “orthologs” function of the babelgene package (Version: 22.9). B cells were subset from this data by the ImmGen predicted cell types from the above process. B cells were scored for cell cycle phase using the same process as described above. The subset B cells were also clustered using a two-step hierarchical clustering approach. The first step of this process was to use the FindClusters function of Seurat 5.1.0^63^ at a high resolution (chosen resolution was 2). The aggregate expression of these 46 clusters was calculated using the AggregateExpression function of Seurat 5.1.0. These clustered cells were then assigned to 10 aggregate clusters using the hclust and cutree functions of the stats package installed in the base R environment (Version: 4.4.3). A principal component analysis to demonstrate the relative expression of the B cells from the individual samples was conducted using the AggregateExpression function of Seurat 5.1.0 to calculate the aggregate expression of each sample before loading the resultant counts matrix into the DESeq2 package (Version: 1.42.1) and transforming the data using the varianceStabilizingTransformation function of DESeq2. Pseudobulk differential gene expression of select B cell clusters was performed by first subsetting the data for the clusters of interest and then calculating the aggregate expression of the individual samples as described in the previous paragraph. Differentially expressed genes were identified using the results function of DESeq2 and setting the contrast to the breast cancer bone metastasis samples compared to the bone marrow control samples. The significance threshold for these differentially expressed genes was a log2 fold change > +/- 1.5 and a p adjusted value < 0.05.

### Statistical analysis

Statistical analysis was performed using Prism 10. Statistical details of the experiments can be found in figure legends. All results are representative of at least three independent experiments.

## REFERENCES

1. Boire, A., Burke, K., Cox, T.R., Guise, T., Jamal-Hanjani, M., Janowitz, T., Kaplan, R., Lee, R., Swanton, C., Vander Heiden, M.G., and Sahai, E. (2024). Why do patients with cancer die? Nat Rev Cancer 24, 578–589. 10.1038/s41568-024-00708-4.

2. Swanton, C., Bernard, E., Abbosh, C., Andre, F., Auwerx, J., Balmain, A., Bar-Sagi, D., Bernards, R., Bullman, S., DeGregori, J., et al. (2024). Embracing cancer complexity: Hallmarks of systemic disease. Cell 187, 1589–1616. 10.1016/j.cell.2024.02.009.

3. Gerstberger, S., Jiang, Q., and Ganesh, K. (2023). Metastasis. Cell 186, 1564–1579. 10.1016/j.cell.2023.03.003.

4. Reagan, M.R., and Rosen, C.J. (2016). Navigating the bone marrow niche: translational insights and cancer-driven dysfunction. Nat Rev Rheumatol 12, 154–168. 10.1038/nrrheum.2015.160.

5. Coleman, R.E., Croucher, P.I., Padhani, A.R., Clezardin, P., Chow, E., Fallon, M., Guise, T., Colangeli, S., Capanna, R., and Costa, L. (2020). Bone metastases. Nat Rev Dis Primers 6, 83. 10.1038/s41572-020-00216-3.

6. Lucas, D. (2021). Structural organization of the bone marrow and its role in hematopoiesis. Curr Opin Hematol 28, 36–42. 10.1097/MOH.0000000000000621.

7. Jackett, K.N., Browne, A.T., Aber, E.R., Clements, M., and Kaplan, R.N. (2024). How the bone microenvironment shapes the pre-metastatic niche and metastasis. Nat Cancer 5, 1800–1814. 10.1038/s43018-024-00854-6.

8. Kaplan, R.N., Riba, R.D., Zacharoulis, S., Bramley, A.H., Vincent, L., Costa, C., MacDonald, D.D., Jin, D.K., Shido, K., Kerns, S.A., et al. (2005). VEGFR1-positive haematopoietic bone marrow progenitors initiate the pre-metastatic niche. Nature 438, 820–827. 10.1038/nature04186.

9. Garner, H., Martinovic, M., Liu, N.Q., Bakker, N.A.M., Velilla, I.Q., Hau, C.S., Vrijland, K., Kaldenbach, D., Kok, M., de Wit, E., and de Visser, K.E. (2025). Understanding and reversing mammary tumor-driven reprogramming of myelopoiesis to reduce metastatic spread. Cancer Cell 43, 1279–1295 e1279. 10.1016/j.ccell.2025.04.007.

10. LaMarche, N.M., Hegde, S., Park, M.D., Maier, B.B., Troncoso, L., Le Berichel, J., Hamon, P., Belabed, M., Mattiuz, R., Hennequin, C., et al. (2024). An IL-4 signalling axis in bone marrow drives pro-tumorigenic myelopoiesis. Nature 625, 166–174. 10.1038/s41586-023-06797-9.

11. Gerber-Ferder, Y., Cosgrove, J., Duperray-Susini, A., Missolo-Koussou, Y., Dubois, M., Stepaniuk, K., Pereira-Abrantes, M., Sedlik, C., Lameiras, S., Baulande, S., et al. (2023). Breast cancer remotely imposes a myeloid bias on haematopoietic stem cells by reprogramming the bone marrow niche. Nat Cell Biol 25, 1736–1745. 10.1038/s41556-023-01291-w.

12. Wculek, S.K., and Malanchi, I. (2015). Neutrophils support lung colonization of metastasis-initiating breast cancer cells. Nature 528, 413–417. 10.1038/nature16140.

13. Coffelt, S.B., Kersten, K., Doornebal, C.W., Weiden, J., Vrijland, K., Hau, C.S., Verstegen, N.J.M., Ciampricotti, M., Hawinkels, L., Jonkers, J., and de Visser, K.E. (2015). IL-17-producing gammadelta T cells and neutrophils conspire to promote breast cancer metastasis. Nature 522, 345–348. 10.1038/nature14282.

14. Li, F., Du, X., Lan, F., Li, N., Zhang, C., Zhu, C., Wang, X., He, Y., Shao, Z., Chen, H., et al. (2021). Eosinophilic inflammation promotes CCL6-dependent metastatic tumor growth. Sci Adv 7. 10.1126/sciadv.abb5943.

15. Monteran, L., Ershaid, N., Scharff, Y., Zoabi, Y., Sanalla, T., Ding, Y., Pavlovsky, A., Zait, Y., Langer, M., Caller, T., et al. (2024). Combining TIGIT Blockade with MDSC Inhibition Hinders Breast Cancer Bone Metastasis by Activating Antitumor Immunity. Cancer Discov 14, 1252–1275. 10.1158/2159-8290.CD-23-0762.

16. Han, Y., Sarkar, H., Xu, Z., Lopez-Darwin, S., Wei, Y., Hang, X., Liu, F., Tran, K., Wang, W., Miller, J.M., et al. (2025). Tumors hijack macrophages for iron supply to promote bone metastasis and anemia. Cell 188, 6335–6354 e6326. 10.1016/j.cell.2025.08.013.

17. Monteran, L., Ershaid, N., Sabah, I., Fahoum, I., Zait, Y., Shani, O., Cohen, N., Eldar-Boock, A., Satchi-Fainaro, R., and Erez, N. (2020). Bone metastasis is associated with acquisition of mesenchymal phenotype and immune suppression in a model of spontaneous breast cancer metastasis. Sci Rep 10, 13838. 10.1038/s41598-020-70788-3.

18. Canon, J.R., Roudier, M., Bryant, R., Morony, S., Stolina, M., Kostenuik, P.J., and Dougall, W.C. (2008). Inhibition of RANKL blocks skeletal tumor progression and improves survival in a mouse model of breast cancer bone metastasis. Clin Exp Metastasis 25, 119–129. 10.1007/s10585-007-9127-1.

19. Simmons, J.K., Hildreth, B.E., 3rd, Supsavhad, W., Elshafae, S.M., Hassan, B.B., Dirksen, W.P., Toribio, R.E., and Rosol, T.J. (2015). Animal Models of Bone Metastasis. Vet Pathol 52, 827–841. 10.1177/0300985815586223.

20. Yu, Y., Li, K., Peng, Y., Wu, W., Chen, F., Shao, Z., and Zhang, Z. (2023). Animal models of cancer metastasis to the bone. Front Oncol 13, 1165380. 10.3389/fonc.2023.1165380.

21. Hiraga, T., and Ninomiya, T. (2019). Establishment and characterization of a C57BL/6 mouse model of bone metastasis of breast cancer. J Bone Miner Metab 37, 235–242. 10.1007/s00774-018-0927-y.

22. Nobre, A.R., Risson, E., Singh, D.K., Di Martino, J.S., Cheung, J.F., Wang, J., Johnson, J., Russnes, H.G., Bravo-Cordero, J.J., Birbrair, A., et al. (2021). Bone marrow NG2(+)/Nestin(+) mesenchymal stem cells drive DTC dormancy via TGFbeta2. Nat Cancer 2, 327–339. 10.1038/s43018-021-00179-8.

23. Lelekakis, M., Moseley, J.M., Martin, T.J., Hards, D., Williams, E., Ho, P., Lowen, D., Javni, J., Miller, F.R., Slavin, J., and Anderson, R.L. (1999). A novel orthotopic model of breast cancer metastasis to bone. Clin Exp Metastasis 17, 163–170. 10.1023/a:1006689719505.

24. Yip, R.K.H., Rimes, J.S., Capaldo, B.D., Vaillant, F., Mouchemore, K.A., Pal, B., Chen, Y., Surgenor, E., Murphy, A.J., Anderson, R.L., et al. (2021). Mammary tumour cells remodel the bone marrow vascular microenvironment to support metastasis. Nat Commun 12, 6920. 10.1038/s41467-021-26556-6.

25. Johnstone, C.N., Smith, Y.E., Cao, Y., Burrows, A.D., Cross, R.S., Ling, X., Redvers, R.P., Doherty, J.P., Eckhardt, B.L., Natoli, A.L., et al. (2015). Functional and molecular characterisation of EO771.LMB tumours, a new C57BL/6-mouse-derived model of spontaneously metastatic mammary cancer. Dis Model Mech 8, 237–251. 10.1242/dmm.017830.

26. Nagasawa, T. (2006). Microenvironmental niches in the bone marrow required for B-cell development. Nat Rev Immunol 6, 107–116. 10.1038/nri1780.

27. Hardy, R.R., Carmack, C.E., Shinton, S.A., Kemp, J.D., and Hayakawa, K. (1991). Resolution and characterization of pro-B and pre-pro-B cell stages in normal mouse bone marrow. J Exp Med 173, 1213–1225. 10.1084/jem.173.5.1213.

28. Heng, T.S., Painter, M.W., and Immunological Genome Project, C. (2008). The Immunological Genome Project: networks of gene expression in immune cells. Nat Immunol 9, 1091–1094. 10.1038/ni1008-1091.

29. Lee, R.D., Munro, S.A., Knutson, T.P., LaRue, R.S., Heltemes-Harris, L.M., and Farrar, M.A. (2021). Single-cell analysis identifies dynamic gene expression networks that govern B cell development and transformation. Nat Commun 12, 6843. 10.1038/s41467-021-27232-5.

30. Seo, S.S., Louros, S.R., Anstey, N., Gonzalez-Lozano, M.A., Harper, C.B., Verity, N.C., Dando, O., Thomson, S.R., Darnell, J.C., Kind, P.C., et al. (2022). Excess ribosomal protein production unbalances translation in a model of Fragile X Syndrome. Nat Commun 13, 3236. 10.1038/s41467-022-30979-0.

31. Gerashchenko, M.V., Nesterchuk, M.V., Smekalova, E.M., Paulo, J.A., Kowalski, P.S., Akulich, K.A., Bogorad, R., Dmitriev, S.E., Gygi, S., Zatsepin, T., et al. (2020). Translation elongation factor 2 depletion by siRNA in mouse liver leads to mTOR-independent translational upregulation of ribosomal protein genes. Sci Rep 10, 15473. 10.1038/s41598-020-72399-4.

32. Liu, F., Ding, Y., Xu, Z., Hao, X., Pan, T., Miles, G., Wang, S., Wu, Y.H., Liu, J., Bado, I.L., et al. (2025). Single-cell profiling of bone metastasis ecosystems from multiple cancer types reveals convergent and divergent mechanisms of bone colonization. Cell Genom 5, 100888. 10.1016/j.xgen.2025.100888.

33. Kitamura, D., Roes, J., Kuhn, R., and Rajewsky, K. (1991). A B cell-deficient mouse by targeted disruption of the membrane exon of the immunoglobulin mu chain gene. Nature 350, 423–426. 10.1038/350423a0.

34. Demircik, F., Buch, T., and Waisman, A. (2013). Efficient B cell depletion via diphtheria toxin in CD19-Cre/iDTR mice. PLoS One 8, e60643. 10.1371/journal.pone.0060643.

35. Hiam-Galvez, K.J., Allen, B.M., and Spitzer, M.H. (2021). Systemic immunity in cancer. Nat Rev Cancer 21, 345–359. 10.1038/s41568-021-00347-z.

36. Chen, F., Han, Y., and Kang, Y. (2021). Bone marrow niches in the regulation of bone metastasis. Br J Cancer 124, 1912–1920. 10.1038/s41416-021-01329-6.

37. Winkler, I.G., Bendall, L.J., Forristal, C.E., Helwani, F., Nowlan, B., Barbier, V., Shen, Y., Cisterne, A., Sedger, L.M., and Levesque, J.P. (2013). B-lymphopoiesis is stopped by mobilizing doses of G-CSF and is rescued by overexpression of the anti-apoptotic protein Bcl2. Haematologica 98, 325–333. 10.3324/haematol.2012.069260.

38. Wu, Q., Zhang, J., Kumar, S., Shen, S., Kincaid, M., Johnson, C.B., Zhang, Y.S., Turcotte, R., Alt, C., Ito, K., et al. (2024). Resilient anatomy and local plasticity of naive and stress haematopoiesis. Nature 627, 839–846. 10.1038/s41586-024-07186-6.

39. Eyles, J.L., Roberts, A.W., Metcalf, D., and Wicks, I.P. (2006). Granulocyte colony-stimulating factor and neutrophils--forgotten mediators of inflammatory disease. Nat Clin Pract Rheumatol 2, 500–510. 10.1038/ncprheum0291.

40. Welte, T., Kim, I.S., Tian, L., Gao, X., Wang, H., Li, J., Holdman, X.B., Herschkowitz, J.I., Pond, A., Xie, G., et al. (2016). Oncogenic mTOR signalling recruits myeloid-derived suppressor cells to promote tumour initiation. Nat Cell Biol 18, 632–644. 10.1038/ncb3355.

41. Carvalho, E., Hugo de Almeida, V., Rondon, A.M.R., Possik, P.A., Viola, J.P.B., and Monteiro, R.Q. (2018). Protease-activated receptor 2 (PAR2) upregulates granulocytecolony stimulating factor (G-CSF) expression in breast cancer cells. Biochem Biophys Res Commun 504, 270–276. 10.1016/j.bbrc.2018.08.169.

42. Loder, F., Mutschler, B., Ray, R.J., Paige, C.J., Sideras, P., Torres, R., Lamers, M.C., and Carsetti, R. (1999). B cell development in the spleen takes place in discrete steps and is determined by the quality of B cell receptor-derived signals. J Exp Med 190, 75–89. 10.1084/jem.190.1.75.

43. Verheijen, M., Rane, S., Pearson, C., Yates, A.J., and Seddon, B. (2020). Fate Mapping Quantifies the Dynamics of B Cell Development and Activation throughout Life. Cell Rep 33, 108376. 10.1016/j.celrep.2020.108376.

44. Cyster, J.G., and Allen, C.D.C. (2019). B Cell Responses: Cell Interaction Dynamics and Decisions. Cell 177, 524–540. 10.1016/j.cell.2019.03.016.

45. Yang, Y., Chen, X., Pan, J., Ning, H., Zhang, Y., Bo, Y., Ren, X., Li, J., Qin, S., Wang, D., et al. (2024). Pan-cancer single-cell dissection reveals phenotypically distinct B cell subtypes. Cell 187, 4790–4811 e4722. 10.1016/j.cell.2024.06.038.

46. Ma, J., Wu, Y., Ma, L., Yang, X., Zhang, T., Song, G., Li, T., Gao, K., Shen, X., Lin, J., et al. (2024). A blueprint for tumor-infiltrating B cells across human cancers. Science 384, eadj4857. 10.1126/science.adj4857.

47. Xu, Y., Mao, Y., Lv, Y., Tang, W., and Xu, J. (2023). B cells in tumor metastasis: friend or foe? Int J Biol Sci 19, 2382–2393. 10.7150/ijbs.79482.

48. Olkhanud, P.B., Damdinsuren, B., Bodogai, M., Gress, R.E., Sen, R., Wejksza, K., Malchinkhuu, E., Wersto, R.P., and Biragyn, A. (2011). Tumor-evoked regulatory B cells promote breast cancer metastasis by converting resting CD4(+) T cells to T-regulatory cells. Cancer Res 71, 3505–3515. 10.1158/0008-5472.CAN-10-4316.

49. Kowanetz, M., Wu, X., Lee, J., Tan, M., Hagenbeek, T., Qu, X., Yu, L., Ross, J., Korsisaari, N., Cao, T., et al. (2010). Granulocyte-colony stimulating factor promotes lung metastasis through mobilization of Ly6G+Ly6C+ granulocytes. Proc Natl Acad Sci U S A 107, 21248–21255. 10.1073/pnas.1015855107.

50. Li, P., Lu, M., Shi, J., Hua, L., Gong, Z., Li, Q., Shultz, L.D., and Ren, G. (2020). Dual roles of neutrophils in metastatic colonization are governed by the host NK cell status. Nat Commun 11, 4387. 10.1038/s41467-020-18125-0.

51. Mouchemore, K.A., and Anderson, R.L. (2021). Immunomodulatory effects of G-CSF in cancer: Therapeutic implications. Semin Immunol 54, 101512. 10.1016/j.smim.2021.101512.

52. Park, S.D., Saunders, A.S., Reidy, M.A., Bender, D.E., Clifton, S., and Morris, K.T. (2022). A review of granulocyte colony-stimulating factor receptor signaling and regulation with implications for cancer. Front Oncol 12, 932608. 10.3389/fonc.2022.932608.

53. Day, R.B., Bhattacharya, D., Nagasawa, T., and Link, D.C. (2015). Granulocyte colony-stimulating factor reprograms bone marrow stromal cells to actively suppress B lymphopoiesis in mice. Blood 125, 3114–3117. 10.1182/blood-2015-02-629444.

54. You, G., Zhang, M., Bian, Z., Guo, H., Xu, Z., Ni, Y., Lan, Y., Yue, W., Gong, Y., Chang, Y., et al. (2022). Decoding lymphomyeloid divergence and immune hyporesponsiveness in G-CSF-primed human bone marrow by single-cell RNA-seq. Cell Discov 8, 59. 10.1038/s41421-022-00417-y.

55. Clezardin, P., Coleman, R., Puppo, M., Ottewell, P., Bonnelye, E., Paycha, F., Confavreux, C.B., and Holen, I. (2021). Bone metastasis: mechanisms, therapies, and biomarkers. Physiol Rev 101, 797–855. 10.1152/physrev.00012.2019.

56. Cheng, J.N., Jin, Z., Su, C., Jiang, T., Zheng, X., Guo, J., Li, X., Chu, H., Jia, J., Zhou, Q., et al. (2025). Bone metastases diminish extraosseous response to checkpoint blockade immunotherapy through osteopontin-producing osteoclasts. Cancer Cell 43, 1093–1107 e1099. 10.1016/j.ccell.2025.03.036.

57. Joseph, G.J., Johnson, D.B., and Johnson, R.W. (2023). Immune checkpoint inhibitors in bone metastasis: Clinical challenges, toxicities, and mechanisms. J Bone Oncol 43, 100505. 10.1016/j.jbo.2023.100505.

58. Boring, L., Gosling, J., Chensue, S.W., Kunkel, S.L., Farese, R.V., Jr., Broxmeyer, H.E., and Charo, I.F. (1997). Impaired monocyte migration and reduced type 1 (Th1) cytokine responses in C-C chemokine receptor 2 knockout mice. J Clin Invest 100, 2552–2561. 10.1172/JCI119798.

59. Rickert, R.C., Roes, J., and Rajewsky, K. (1997). B lymphocyte-specific, Cre-mediated mutagenesis in mice. Nucleic Acids Res 25, 1317–1318. 10.1093/nar/25.6.1317.

60. Buch, T., Heppner, F.L., Tertilt, C., Heinen, T.J., Kremer, M., Wunderlich, F.T., Jung, S., and Waisman, A. (2005). A Cre-inducible diphtheria toxin receptor mediates cell lineage ablation after toxin administration. Nat Methods 2, 419–426. 10.1038/nmeth762.

61. Holl, T.M., Haynes, B.F., and Kelsoe, G. (2010). Stromal cell independent B cell development in vitro: generation and recovery of autoreactive clones. J Immunol Methods 354, 53–67. 10.1016/j.jim.2010.01.007.

62. Zheng, G.X., Terry, J.M., Belgrader, P., Ryvkin, P., Bent, Z.W., Wilson, R., Ziraldo, S.B., Wheeler, T.D., McDermott, G.P., Zhu, J., et al. (2017). Massively parallel digital transcriptional profiling of single cells. Nat Commun 8, 14049. 10.1038/ncomms14049.

63. Hao, Y., Hao, S., Andersen-Nissen, E., Mauck, W.M., 3rd, Zheng, S., Butler, A., Lee, M.J., Wilk, A.J., Darby, C., Zager, M., et al. (2021). Integrated analysis of multimodal single-cell data. Cell 184, 3573–3587 e3529. 10.1016/j.cell.2021.04.048.

64. Aran, D., Looney, A.P., Liu, L., Wu, E., Fong, V., Hsu, A., Chak, S., Naikawadi, R.P., Wolters, P.J., Abate, A.R., et al. (2019). Reference-based analysis of lung single-cell sequencing reveals a transitional profibrotic macrophage. Nat Immunol 20, 163–172. 10.1038/s41590-018-0276-y.

65. Love, M.I., Huber, W., and Anders, S. (2014). Moderated estimation of fold change and dispersion for RNA-seq data with DESeq2. Genome Biol 15, 550. 10.1186/s13059-014-0550-8.

66. Zhou, Y., Zhou, B., Pache, L., Chang, M., Khodabakhshi, A.H., Tanaseichuk, O., Benner, C., and Chanda, S.K. (2019). Metascape provides a biologist-oriented resource for the analysis of systems-level datasets. Nat Commun 10, 1523. 10.1038/s41467-019-09234-6.

67. Kowalczyk, M.S., Tirosh, I., Heckl, D., Rao, T.N., Dixit, A., Haas, B.J., Schneider, R.K., Wagers, A.J., Ebert, B.L., and Regev, A. (2015). Single-cell RNA-seq reveals changes in cell cycle and differentiation programs upon aging of hematopoietic stem cells. Genome Res 25, 1860–1872. 10.1101/gr.192237.115.

