## Supplemental information for "Bone marrow B cell collapse promotes bone metastasis in breast cancer"

### SUPPLEMENTAL FIGURES

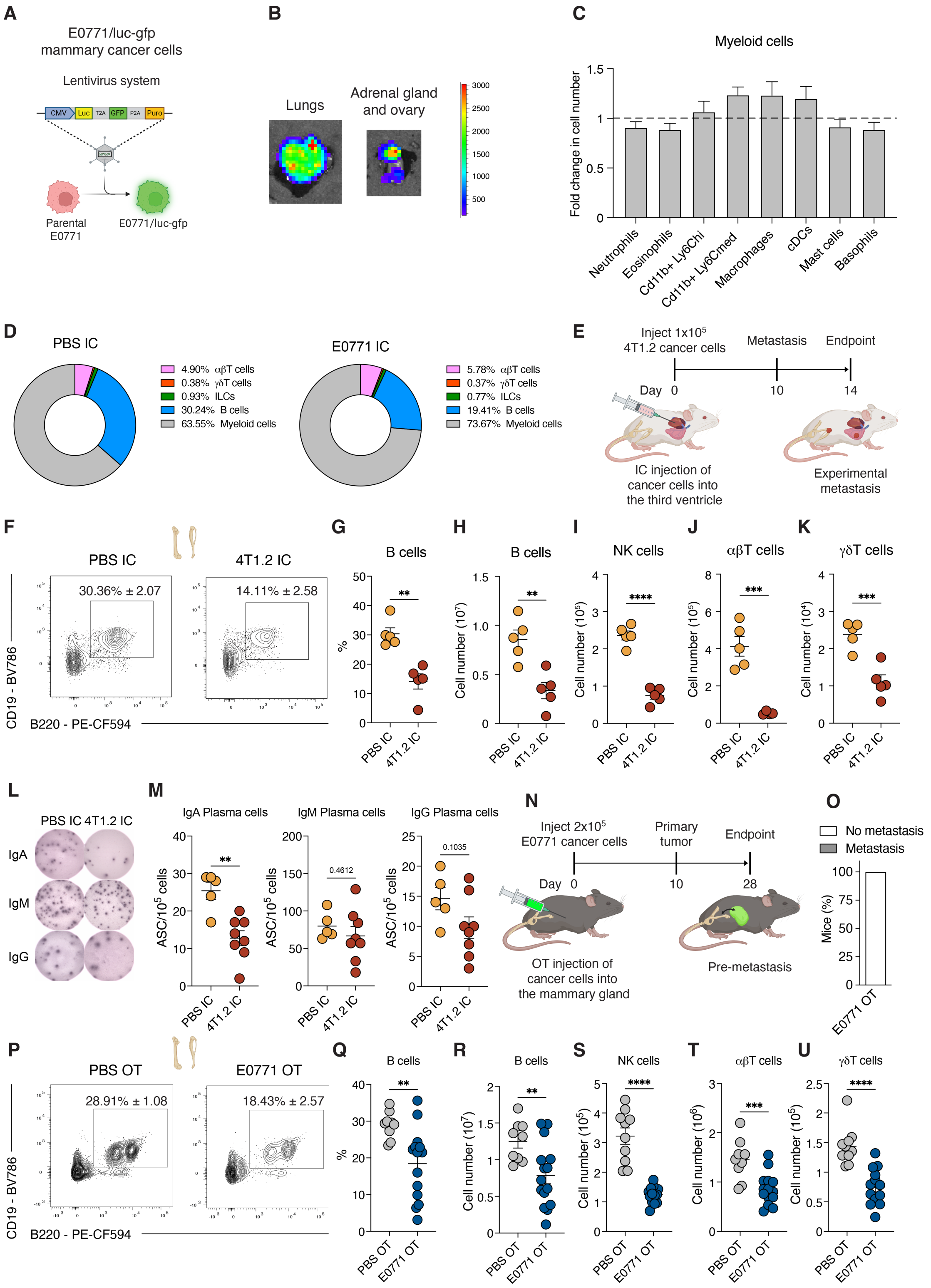


Teijeiro et al. Figure S1

**Figure S1. Mammary tumor cells remodel the bone marrow lymphoid niche, related to Figure 1.**

(A) Schematic representation of the E0771-*gfp-luc* cell line.

(B) Representative images of *ex vivo* bioluminescence in organs from C57BL/6 mice with IC injection of E0771 cells. Color scale represents bioluminescence counts.

(C) Fold changes in absolute numbers of various myeloid cells in bone marrow of C57BL/6 mice injected with E0771 cells IC at day 17 compared to C57BL/6 mice injected with PBS IC.

(D) Pie chart representing the relative abundance of bone marrow immune subsets in C57BL/6 mice injected with PBS or E0771 cells IC at day 17.

(E) Experimental layout. IC injection of 4T1.2 mammary tumor cells is performed in BALB/c mice to induce experimental metastasis.

(F) Representative contour plots of bone marrow $CD45^+$ $TCR\beta^-$ $CD138^-$ $B220^+$ $CD19^+$ B cells in BALB/c mice injected with PBS or 4T1.2 cells IC at day 14.

(G, H) Frequency (G) and absolute numbers (H) of bone marrow $CD45^+$ $TCR\beta^-$ $CD138^-$ $B220^+$ $CD19^+$ B cells in mice described in (F).

(I-K) Absolute numbers of bone marrow $CD45^+$ $CD90.2^+$ $TCR\beta^-$ $NK1.1^+$ NK cells (I), $CD45^+$ $CD90.2^+$ $TCR\beta^+$ $\gamma\delta TCR^-$ $\alpha\beta$T cells (J), and $CD45^+$ $CD90.2^+$ $\gamma\delta TCR^+$ $TCR\beta^-$ $\gamma\delta$T cells (K) in mice described in (F).

(L, M) Representative pictures of ELISPOT wells (L) and quantification (M) of the frequency of antibody secreting cells (ASCs) in bone marrow of mice described in (F).

(N) Experimental layout. Orthotopic (OT) injection of E0771 mammary tumor cells is performed in the mammary gland of C57BL/6 mice. Mice were euthanized 28 days after injection.

(O) Incidence of metastasis in C57BL/6 mice with OT injection of E0771 cells at day 28.

(P) Representative contour plots of bone marrow $CD45^+$ $TCR\beta^-$ $CD138^-$ $B220^+$ $CD19^+$ B cells in C57BL/6 mice injected with PBS or E0771 cells OT at day 28.

(Q, R) Frequency (Q) and absolute numbers (R) of bone marrow CD45$^+$ TCRβ$^-$ CD138$^-$ B220$^+$ CD19$^+$ B cells in mice described in (P).

(S-U) Absolute numbers of bone marrow CD45$^+$ CD90.2$^+$ TCRβ$^-$ NK1.1$^+$ NK cells (S), CD45$^+$ CD90.2$^+$ TCRβ$^+$ γδTCR$^-$ αβT cells (T), and CD45$^+$ CD90.2$^+$ γδTCR$^+$ TCRβ$^-$ γδT cells (U) in mice described in (P).

Statistical analysis was performed using unpaired two-tailed Student's t test (C, G-K, M, Q-U). Data are represented as means ± standard error of the mean (SEM). Each dot represents a mouse in (G-K, M, and Q-U). $**p \leq 0.01$, $***p \leq 0.001$, $****p \leq 0.0001$.

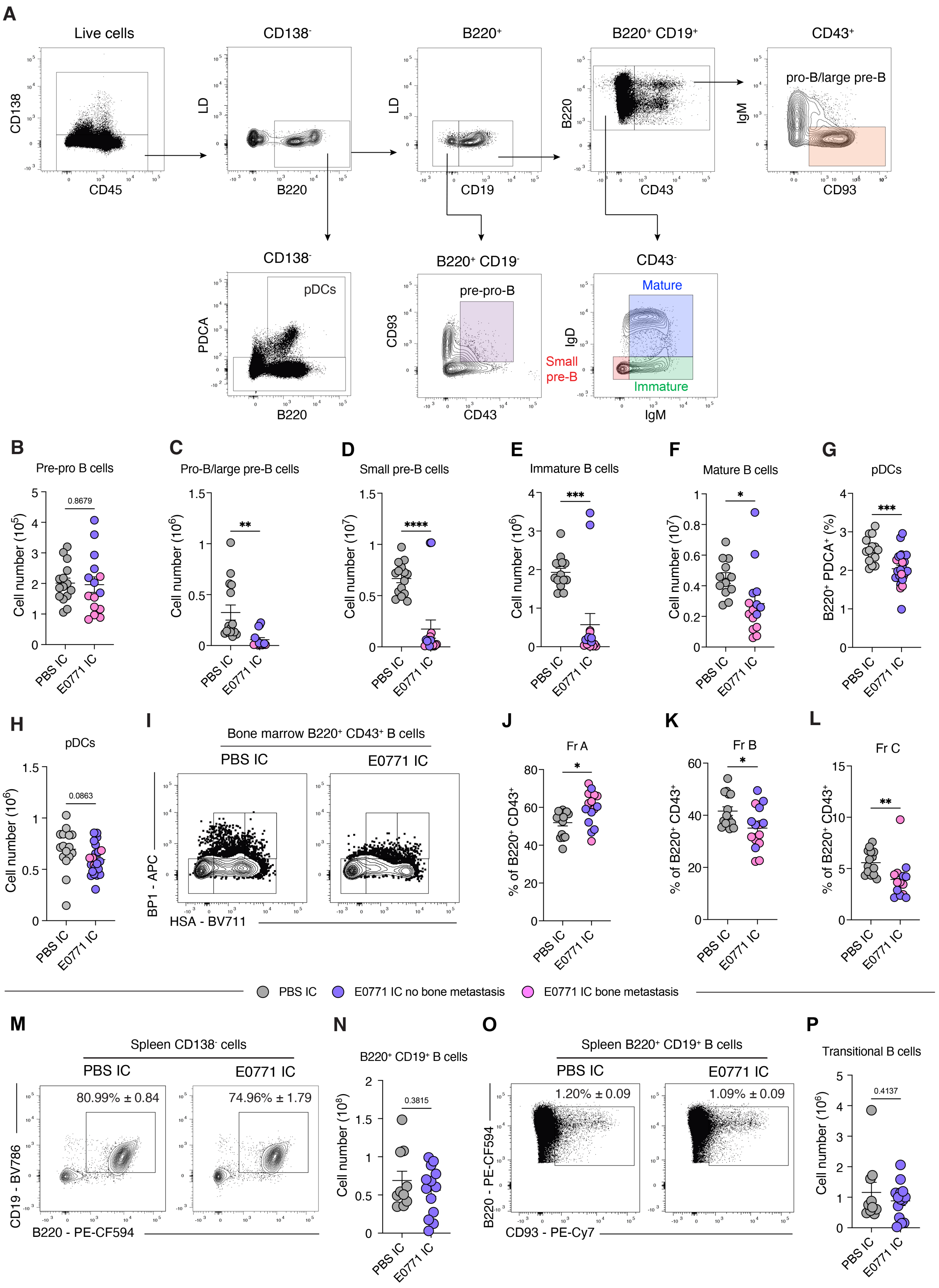


Teijeiro et al. Figure S2

**Figure S2. Mammary tumor cells impair B cell development, related to Figure 2.**

(A) Gating strategy for B cell subsets and pDCs. LD: Live/dead dye.

(B-F) Absolute numbers of bone marrow B cell subsets in C57BL/6 mice with IC injection of PBS or E0771 cells at day 17.

(H, I) Frequency (H) and absolute numbers (I) of B220$^{+}$ PDCA$^{+}$ pDCs in mice described in (B).

(I-L) Representative contour plots (I) and frequency of bone marrow B220$^{+}$ CD43$^{+}$ BP1$^{-}$ HSA$^{-}$ fraction (Fr) A B cells (J), B220$^{+}$ CD43$^{+}$ BP1$^{-}$ HSA$^{+}$ Fr B B cells (K), and B220$^{+}$ CD43$^{+}$ BP1$^{+}$ HSA$^{+}$ Fr C B cells (L) in mice described in (B).

(M, N) Representative contour plots (M) and absolute numbers (N) of splenic B220$^{+}$ CD19$^{+}$ B cells in mice described in (B).

(O, P) Representative dot plots (O) and absolute numbers (P) of splenic B220$^{+}$ CD19$^{+}$ CD93$^{+}$ transitional B cells in mice described in (B).

Statistical analysis was performed using unpaired two-tailed Student's t test (B-H, J-L, N, P). Data are represented as means ± SEM. Each dot represents a mouse in (B-H, J-L, N, P). ∗p ≤ 0.05, ∗∗p ≤ 0.01∗∗∗p ≤ 0.001, ∗∗∗∗p ≤ 0.0001.

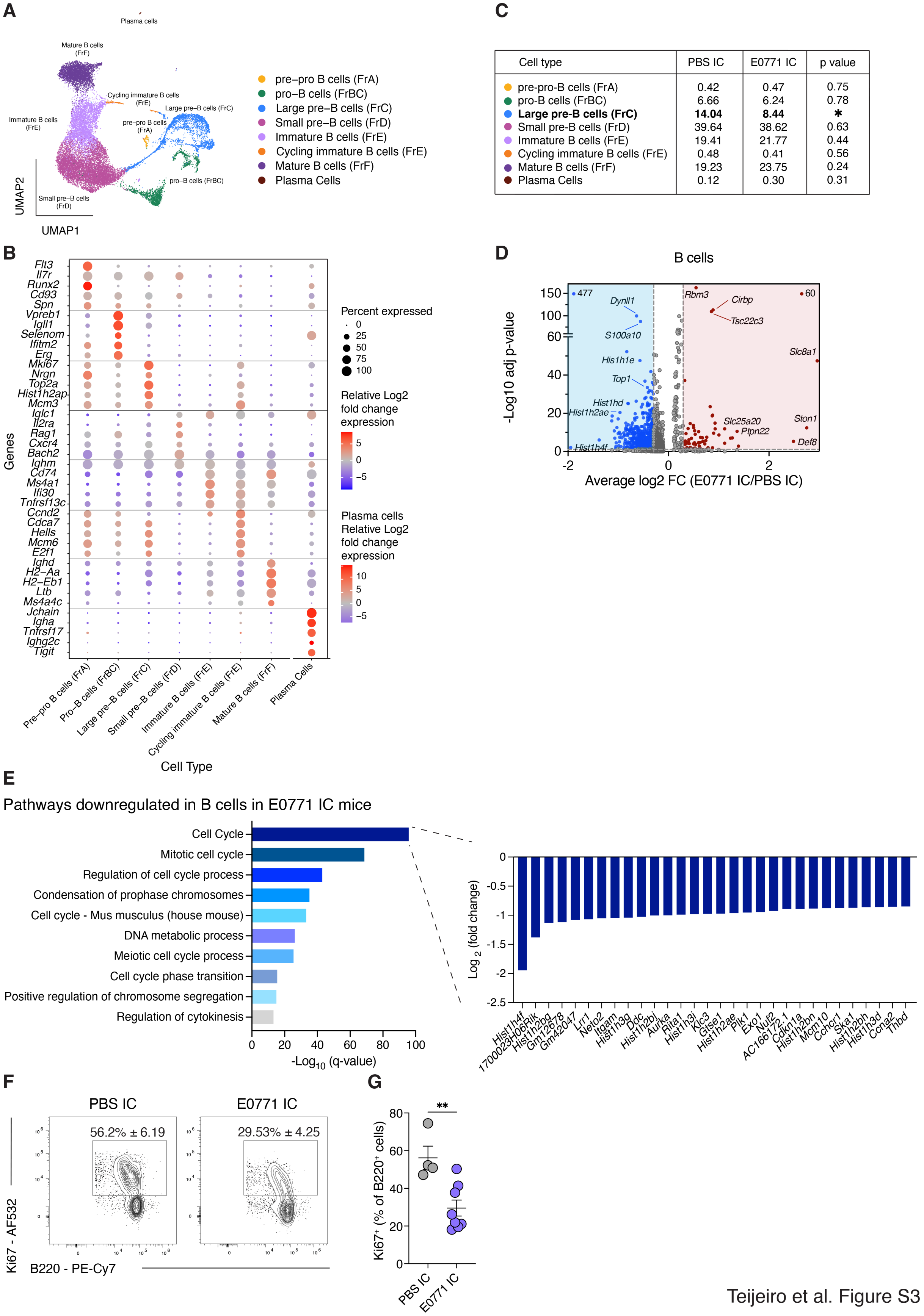


| Cell type | PBS IC | E0771 IC | p value |
|---|---|---|---|
| pre-pro-B cells (FrA) | 0.42 | 0.47 | 0.75 |
| pro-B cells (FrBC) | 6.66 | 6.24 | 0.78 |
| **Large pre-B cells (FrC)** | **14.04** | **8.44** | ***** |
| Small pre-B cells (FrD) | 39.64 | 38.62 | 0.63 |
| Immature B cells (FrE) | 19.41 | 21.77 | 0.44 |
| Cycling immature B cells (FrE) | 0.48 | 0.41 | 0.56 |
| Mature B cells (FrF) | 19.23 | 23.75 | 0.24 |
| Plasma Cells | 0.12 | 0.30 | 0.31 |

Teijeiro et al. Figure S3

**Figure S3. Mammary tumor cells redefine the bone marrow B cell transcriptional landscape, related to Figure 3.**

(A) UMAP projection showing bone marrow B cell clusters of C57BL/6 mice with IC injection of PBS or E0771 cells at day 17 analyzed by scRNAseq and annotated using the ImmGen database.

(B) Dot plot of highly expressed genes in each cluster from B cell scRNAseq analysis, used for cluster annotation.

(C) Frequency of scRNAseq clusters of bone marrow B cells from mice described in (A).

(D) Volcano plot displaying pseudobulk differentially expressed genes in B cells between PBS IC and E0771 IC mice.

(E) Bar graph representing pathways downregulated in B cells from E0771 IC mice relative to PBS IC mice.

(F, G) Representative contour plots (F) and frequency (G) of $Ki67^{+}$ cells in bone marrow $B220^{+}$ B cells from mice described in (A).

Statistical analysis was performed using unpaired two-tailed Student's t test (C, G). Data are represented as means ± SEM. Each dot represents a cell in (A), a gene in (D), and a mouse in (G). $*p \leq 0.05$, $**p \leq 0.01$.

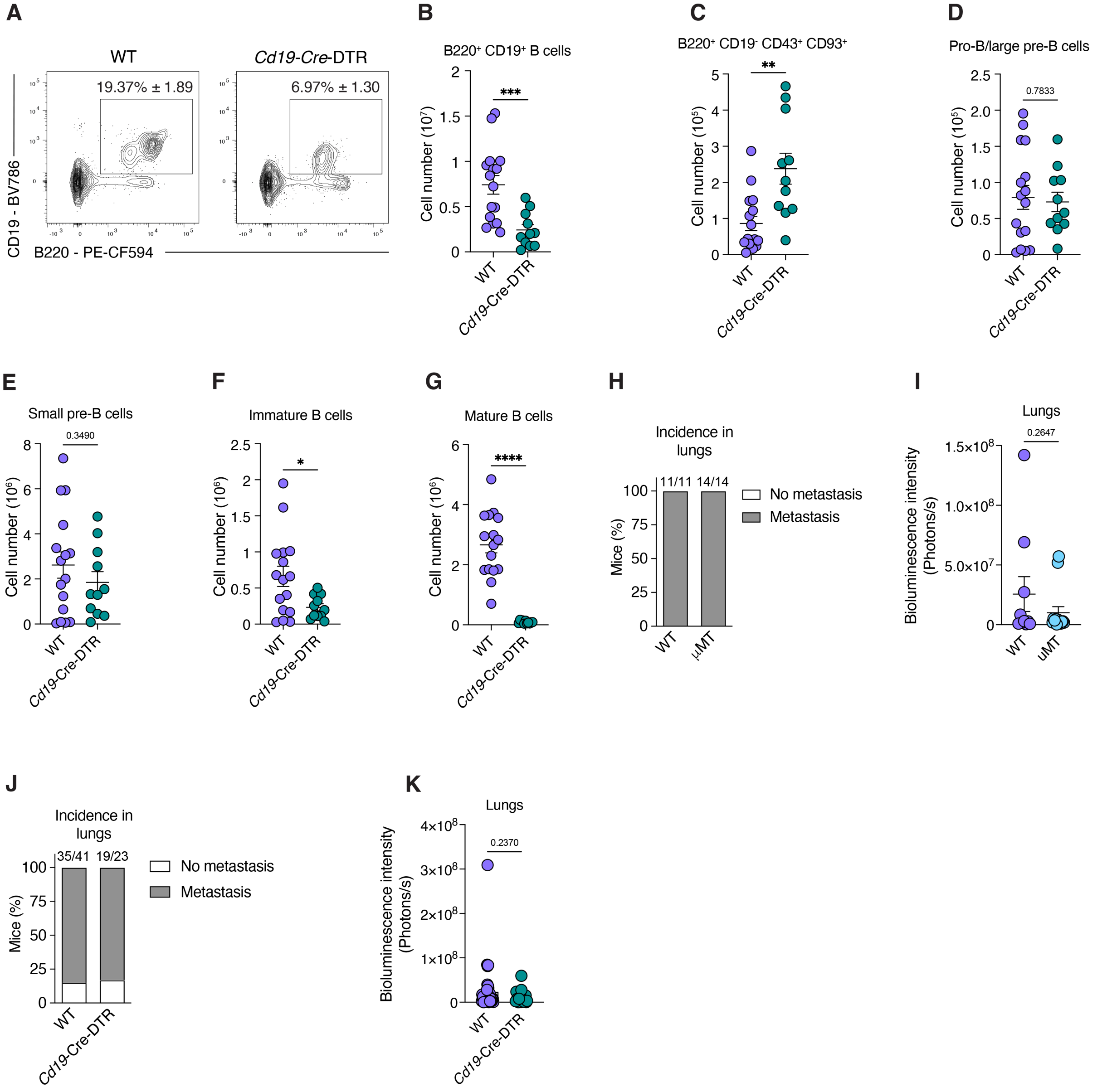


Teijeiro et al. Figure S4

**Figure S4. Experimental B cell loss promotes mammary tumor bone metastasis, related to Figure 5.**

(A, B) Representative contour plots (A) and absolute numbers (B) of bone marrow B cells in wildtype (WT) *Cd19*-Cre-DTR mice injected IC with E0771 mammary tumor cells and analyzed 17 days after IC injection.

(C-G) Absolute numbers of bone marrow B220$^{+}$ CD19$^{-}$ CD43$^{+}$ CD93$^{+}$ cells (C), B220$^{+}$ CD19$^{+}$ CD43$^{+}$ CD93$^{+}$ IgM$^{-}$ pro-B/large pre-B cells (D), B220$^{+}$ CD19$^{+}$ CD43$^{-}$ IgM$^{-}$ IgD$^{-}$ small pre-B cells (E), B220$^{+}$ CD19$^{+}$ CD43$^{-}$ IgM$^{+}$ IgD$^{-}$ immature B cells (F), and B220$^{+}$ CD19$^{+}$ CD43$^{-}$ IgM$^{+}$ IgD$^{+}$ mature B cells (G) in mice described in (A).

(H, I) Lung metastasis incidence (H) and quantification of luciferase signal in lungs with mammary tumor metastasis (I) in WT and μMT mice injected IC with E0771 mammary tumor cells and analyzed 17 days after injection.

(J, K) Lung metastasis incidence (J) and quantification of luciferase signal in lungs with mammary tumor metastasis (K) in mice described in (A).

Statistical analysis was performed using unpaired two-tailed Student's t test (B-G, I, K) or Fisher's exact test (H, J). Data are represented as means ± SEM. Each dot represents a mouse in (B-G) and an organ in (I, K). $*p \leq 0.05$, $**p \leq 0.01$, $***p \leq 0.001$, $****p \leq 0.0001$.

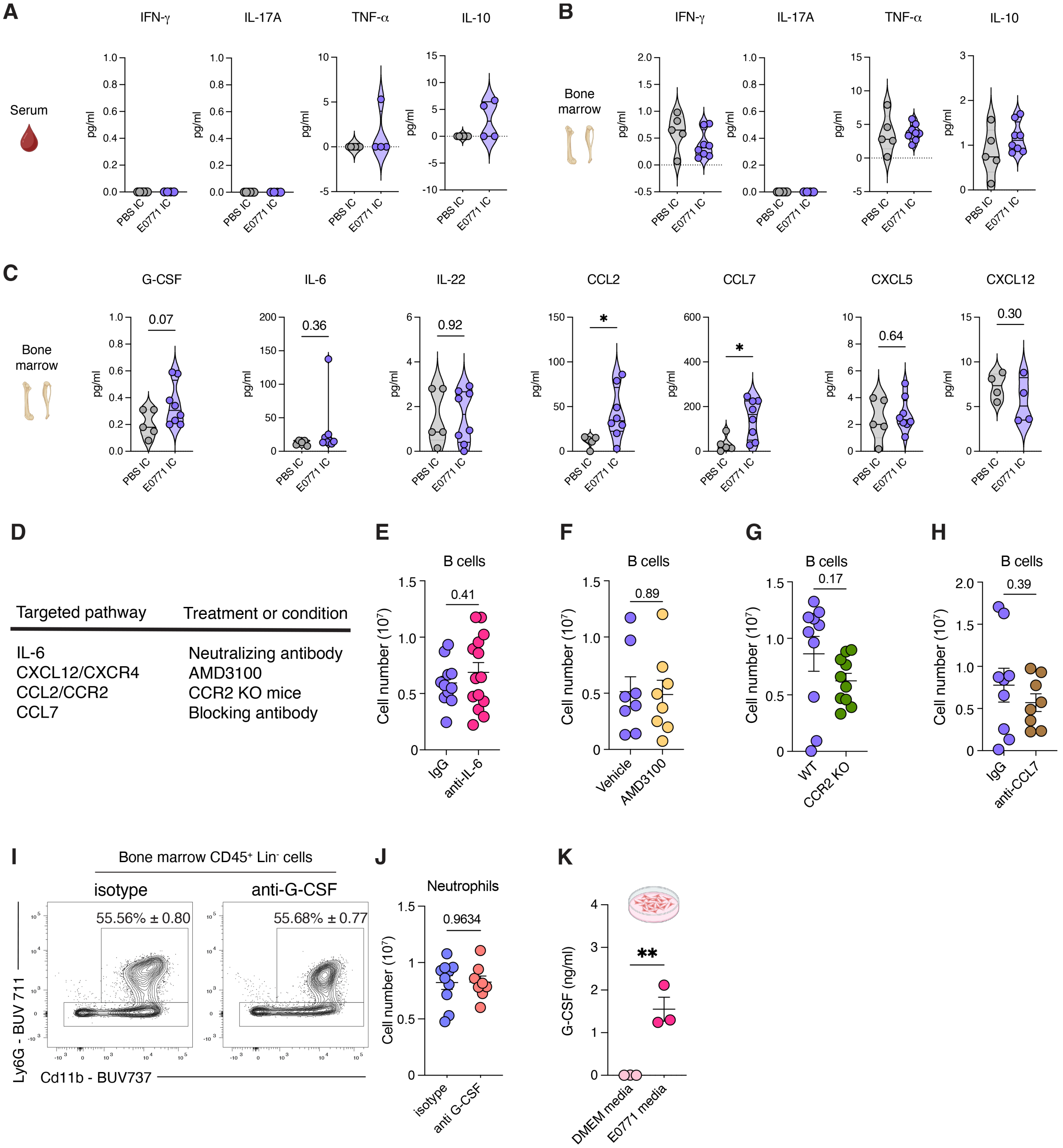


Teijeiro et al. Figure S5

**Figure S5. G-CSF is required for B cell loss and mammary tumor bone metastasis, related to Figure 6.**

(A-C) Cytokine and chemokine levels in blood (A) and bone marrow lysates (B, C) of C57BL/6 mice with IC injection of PBS or E0771 cells at day 17.

(D) Table summarizing the *in vivo* screening of potential cytokines and chemokines mediating B cell loss in E0771 IC mice.

(E-H) Absolute numbers of bone marrow B220$^+$ CD19$^+$ bone marrow B cells in E0771 IC mice treated with anti-IL-6 (E), treated with AMD3100 (F), CCR2 KO mice (G), and treated with anti-CCL7 (H) and analyzed 17 days after tumor cell injection.

(I, J) Representative contour plots (I) and absolute numbers (J) of CD45$^+$ Lineage (Lin = TCRβ, γδTCR, B220, NK1.1)$^-$ Cd11b$^+$ LyG$^+$ neutrophils in bone marrow from C57BL/6 mice treated with G-CSF antibody or the isotype control and injected IC with E0771 mammary tumor cells and analyzed at day 15.

(K) G-CSF levels in DMEM media or media from cultured E0771 cells after 48 hours of incubation.

Statistical analysis was performed using unpaired two-tailed Student's t test (A-C, E-H, J, K). Data are represented as means ± SEM. Each dot represents a mouse in (A-C, E-H, and J) and a cell culture sample in (K). $*p \leq 0.05$, $**p \leq 0.01$.
